# Ezh2 Delays Activation of Differentiation Genes During Normal Cerebellar Granule Neuron Development and in Medulloblastoma

**DOI:** 10.1101/2024.11.21.624171

**Authors:** James Purzner, Alexander S. Brown, Teresa Purzner, Lauren Ellis, Sara Broski, Ulrike Litzenburger, Kaytlin Andrews, Aryaman Sharma, Xin Wang, Michael D. Taylor, Yoon-Jae Cho, Margaret T. Fuller, Matthew P. Scott

## Abstract

Medulloblastoma (MB) is the most common malignant brain tumour in children. The Sonic Hedgehog (SHH)-medulloblastoma subtype arises from the cerebellar granule neuron lineage. Terminally differentiated neurons are incapable of undergoing further cell division, so an effective treatment for this tumour could be to force neuronal differentiation. Differentiation therapy provides a potential alternative for patients with medulloblastoma who harbor mutations that impair cell death pathways (TP53), which is associated a with high mortality. To this end, our goal was to explore epigenetic regulation of cerebellar granule neuron differentiation in medulloblastoma cells. Key regulators were discovered using chromatin immunoprecipitation with high-throughput sequencing. DNA-bound protein and chromatin protein modifications were investigated across all genes. We discovered that Ezh2-mediated tri-methylation of the H3 histone (H3K27me3), occurred on more than half of the 787 genes whose transcription normally increases as granule neurons terminally differentiate. Conditional knockout of *Ezh2* led to early initiation of differentiation in granule neuron precursors (GNPs), but only after cell cycle exit had occurred. Similarly, in MB cells, neuronal differentiation could be induced by preventing H3K27me3 modifications using an Ezh2 inhibitor (UNC1999), but only when UNC1999 was combined with forced cell cycle exit driven by a CDK4/6 inhibitor (Palbociclib). Ezh2 emerges as a powerful restraint upon post-mitotic differentiation during normal GNP development and combination of Ezh2 inhibition with cell cycle exit leads to MB cell differentiation.

## Introduction

The Hedgehog (Hh) pathway is one of the major regulators of cerebellar granule neuron precursor (GNP) proliferation. Sonic hedgehog (Shh) protein secreted from Purkinje neurons drives proliferation of GNPs (*1–3*) through transcriptional regulation of *Cyclin D1* and *N-myc* (*4, 5*) among other targets. Despite ongoing exposure to Shh, GNPs leave the cell cycle and undergo timed and reliable differentiation, a switch that occurs correctly about 50 billion times in the developing human brain. When differentiation and neurogenesis fail, for example due to Shh pathway mutations that lead to unrestrained Shh target gene activation, GNPs continue to proliferate and may form medulloblastomas (MBs). Yet even in genetic models of MB that are 100% penetrant, where Shh target genes are dramatically activated, the vast majority of GNPs still differentiate into functional granule neurons (GNs) (*6*). Thus, regulation that promotes differentiation has the potential to overcome a potent mitogenic signal.

GNPs are the cell type of origin for the Shh subtype of MBs, which make up a quarter to a third of MBs. Children with Shh MB are treated intensively with surgical resection followed by craniospinal radiation, one year of multi-drug chemotherapy, and targeted inhibition of the Hh pathway using Smo inhibitors. Unfortunately, in patients with germline Hh pathway mutations, or intratumoral mutations in *P53*, the five-year overall survival is halved, from 81% to 41% (*7, 8*). Making matters worse, patients with *P53* mutations often have amplifications in downstream targets like *GLI2* and *NMYC* (*9, 10*), meaning that they will not respond to inhibitors that target the SMO protein.

An alternative therapeutic approach is to force tumor cells into terminal differentiation, which can be less toxic than alkylating chemotherapy or radiation (*11–13*) and does not rely on apoptosis or other cell death machinery. The optimal procedure for inducing terminal differentiation is likely to vary from cell type to cell type, since differentiation of neurons depends on different genes than, for example, differentiating blood cells. Neuronal differentiation of MB does occur spontaneously, as it has been commonly observed in human MB pathology specimens for over 30 years (*14–16*). BMP4 induces differentiation of MB cells, reflecting its normal effect on GNPs (*17, 18*). Unfortunately, no pharmaceutical BMP agonist currently exists. Developmental regulation of cerebellar granule neurons potentially provides us with blueprints for engineering a differentiation therapy that is specific to Shh-subtype MB differentiation.

GNPs undergo approximately eight symmetric divisions (*19*), during transit amplification in the external granule layer (EGL) (Fig 1A). As GNPs divide they travel inward from the pial surface, eventually exiting the cell cycle. GNP movement, morphogenesis, and synapse formation lasts for about 10 days after cell cycle exit (*20–22*). Following the last mitosis the cells remain in the inner EGL (iEGL) for approximately 48 hours. While in the iEGL the GNs begin to extend two lateral processes, then a deep process (Fig 1B). Each GN nucleus migrates inward through the deep process, along Bergmann glia and past the Purkinje cell layer, into the Internal Granule Layer (IGL). The final shape of a GN is a T, with most of the cell’s three processes located in the molecular layer and the nucleus in the IGL. For each step of GNP differentiation, positive and negative regulators have been discovered (*23, 24*). The extensive movement and morphological changes involve the changing expression of hundreds of genes, as shown by recent transcriptional profiling (*25*).

**Figure 1:**
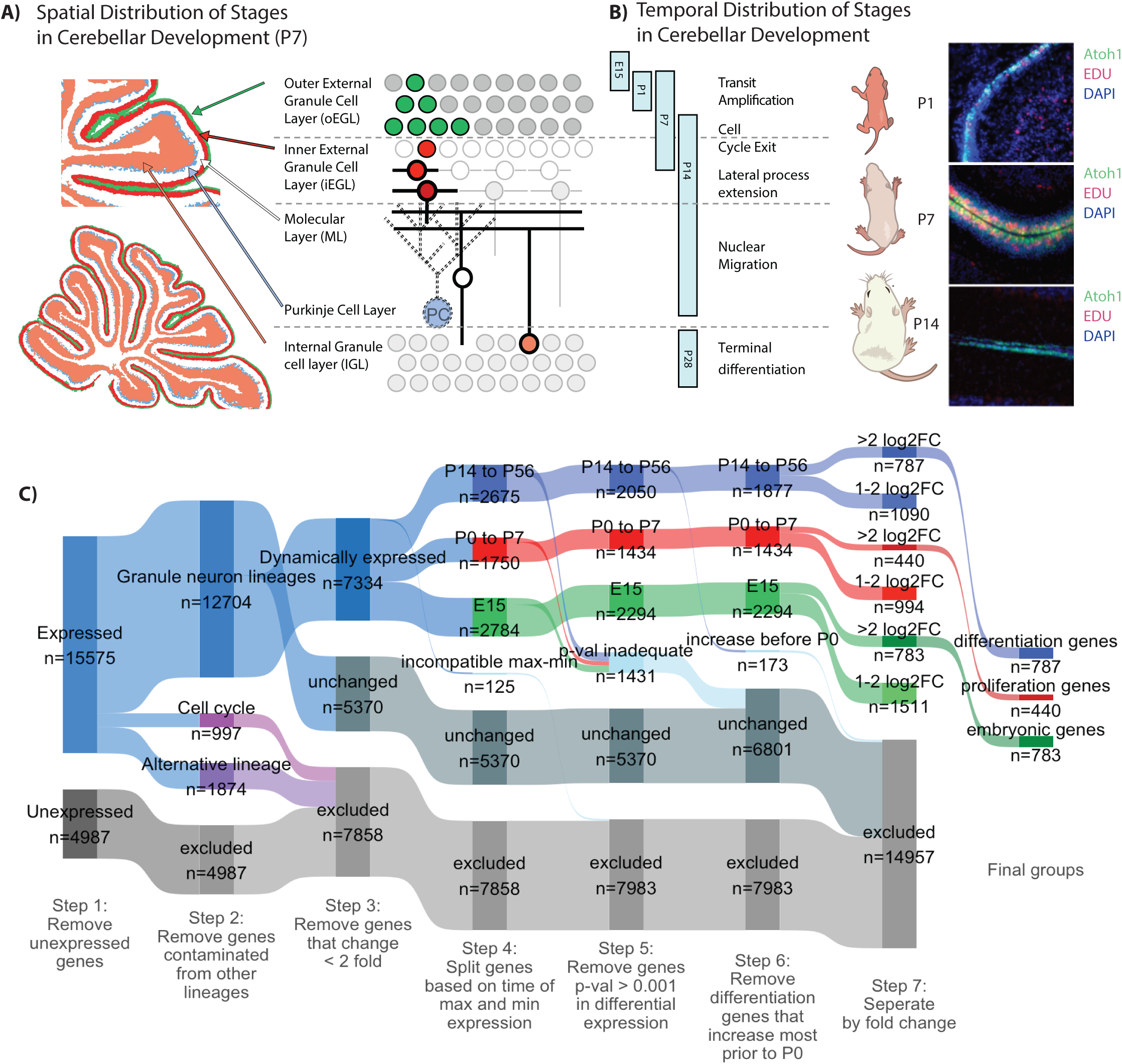
Identifying a high confidence group of genes that increase with GNP differentiation. **A)** Schematic representation of the spatial distribution of stages of GNP development represented in the P7 mouse cerebellum. B) Schematic representation of the temporal distribution of the stages of GNP development. Stages represented in the RNASeq data are indicated by vertical bars. Representative sagittal slices of the cerebellum from mice harboring a *Math1-GFP* reporter, injected with EDU (1.5 h, red) and DAPI (blue) at P1, P7, and P14. **C**) A Sankey diagram describes how genes are partitioned in order to identify high confidence differentiation genes. Briefly, genes are segregated by the following steps: 1, excluded genes by non-expression; 2, excluded genes by contamination from non granule neuron lineage identified through scRNAseq; 3, excluded by minimal change in expression during development; 4, segregated genes by the time of minimum and maximum expression (highest P14 to P6 differentiation genes, highest P0-7 proliferation genes, highest at E15 embryonic genes); 5 excluded genes by low statistical confidence; 6, exclude genes that increase prior to P0; 7, separate by fold-change (>2 fold log2 fold change or 1-2 log2 fold change).

We set out to explore epigenetic regulation of genes whose expression increases during differentiation. Many genes whose transcription change during differentiation encode chromatin-remodeling complexes, implying coordination at the level of transcription (*25*). In addition, SHH MB harbor somatic mutations in genes which impact a number chromatin modifying complexes including genes that impact H3 methylation at K27 and K4 (*26*). A mechanism that represses pro-differentiation genes potentially could be inhibited by an anti-cancer drug and would potentially stimulate differentiation of MBs.

Our plan was to comprehensively describe histone modifications and critical transcription factors at promoters and enhancers of genes that become active during GNP development, to discover inhibitors of differentiation. We found that more than half of the GNP differentiation genes are associated with histones modified to be H3K27me3 and H3K4me3. H3K27 methylation is carried out by the Enhancer of zeste homolog 2 (EZH2) histone methyltransferase, a component of one of the Polycomb Repressive Complexes (PRC2) that were discovered in Drosophila (*27, 28*). H3K4me3 is formed by the MLL complex (*29*) and was not further investigated in our studies. The frequency of H3K27 modifications suggested that inhibition or mutation of EZH2 could lead to premature differentiation, and perhaps to differentiation and growth cessation by MB cells. In *Ezh2* mouse mutants we observed accelerated differentiation of GNPs that had exited the cell cycle. We were able to induce differentiation of MB cells by combining an EZH2 inhibitor with a CDK4/6 inhibitor that arrested cells in G0.

## Results

### Identification of genes activated at the onset of GNP differentiation

We used Percoll fractionation to enrich for GNPs from wild-type mouse cerebella at three key stages, proliferation initiation (P1), peak proliferation (P7), and the onset of differentiation (P14), and then extracted RNA (Materials and Methods; the RNA preparations used are diagrammed in blue in Fig. 1C). We purified earlier-stage GNPs from E15.5 mouse embryos; these had seen little or no Shh (Materials and Methods). E15.5-stage GNPs were also isolated by fluorescence-activated cell sorting (FACS) from E15.5 mice harboring a *Math1-GFP* reporter (*30*). *Math1-GFP* is expressed in neural progenitor cells from the dorsal neural tube, adjacent to the roof plate, that are committed to a GNP lineage. The reporter remains active until GNPs complete their final mitosis.

Data for GNPs from stages later than P14 were obtained using the Translating Ribosome Affinity Purification (TRAP) dataset made available by the Hatten lab (*25*). TRAP data were obtained by purifying mRNA from post-proliferative stages of GNP differentiation (P18, P28) in cells expressing *NeuroD1,* which is transcribed as GNs exit from the cell cycle. TRAP has the benefit of allowing detection and quantification of RNA in the delicate granule neuron processes which are otherwise lost when cerebellar tissue is broken up. *NeuroD1* is not expressed at E15, so earlier time points were not accessible using TRAP. Our RNAseq data were combined with the NeuroD1-TRAP data (23) to achieve a picture of transcription from the earliest stages of GNP formation at E15 to their terminal state at P56.

To identify genes that increase transcription during differentiation we followed a multistep process (Fig. 1C). First, genes were excluded if their transcript level was below a cutoff, established using the maximum expression time point of each dataset in both our RNAseq and the Hatten lab TRAP data, and fitting a mixture of two probability distributions to the bimodal data (expressed and unexpressed). Next, we excluded genes that were potentially contaminants from other cell types. Percoll-fractionated samples from P7 mouse cerebellum typically contain about 95% GNPs and 5% contaminating cells (Fig. 1C, blue v. gray). Genes that are highly expressed in non-GNP cerebellar cells, and scarcely expressed in GNPs, were identified and excluded based upon single-cell RNAseq (scRNAseq) of the developing mouse cerebellum from E10 to P14 (*31*). 1874 genes that were represented by transcripts in bulk cerebellar RNA were determined to be likely contaminants based upon the results of the scRNAseq. Third, genes that vary substantially during the cell cycle were removed as well.

GNP differentiation genes were grouped according to when during development they had maximum transcript abundance. Genes whose transcript levels changed less than 2-fold between developmental stages were categorized as unchanged.

2050 genes were most highly represented in RNA between P14 and P56 and were categorized as differentiation genes. Some peaked early in differentiation, and some peaked during maximal GNP proliferation from P1 to P7. We used the time at which a gene’s transcript reached 50% of its maximum (t50) to describe when genes were undergoing the most transcriptional changes (Fig S1A-C). If a gene reached 50% of its maximum transcript level before P0 it was set aside. Our focus on differentiation therapy means we are primarily interested in genes that increase after the period of peak proliferation, as this is most similar to MB.

787 genes increased 4-fold to 12.4 fold at or after P14, comparing the lowest to highest contiguous time points, and were categorized as differentiation genes (Fig 1C and Supplemental Table 1). Many of the differentiation genes are involved in neuronal function with 181 genes impacting membrane potential or synaptic signalling, which includes 45 ion channels and 45 transporters. Genes involved in cell morphogenesis make up another 57 genes including cell adhesion proteins and another 28 proteins involved in the cytoskeleton. In addition to these terminal effectors of building a functional neuron we also see an increase in 50 transcription factors, which could have extensive downstream impact on transcription (Supplemental Table 2).

Transcripts of a different set of 440 genes were highest (>= 4 fold) during proliferative stages P0-P7. As expected, these proliferation genes included *Gli1,* a direct transcriptional target of the Shh signal transduction pathway that drives GNP proliferation, as well as cell cycle regulators including the G1/S phase cyclin *Ccnd1.* The majority of the genes that have *lower* RNA abundance as GNPs approach cell cycle exit are known regulators or components of the cell cycle such as components of the E2f-Rb complex which controls expression of S-phase cell cycle related genes. Transcript levels for a third set of 783 genes were highest (>= 4-fold change) in the E15 pre-Shh GNPs (*32, 33*).

We checked a putative differentiation gene to see if its protein abundance reflected the observed changes in RNA levels. P7 cerebellar sections (Fig S1E) and GNPs differentiated *in vitro* (Fig S1F,G) had changes in protein abundance that agree with the RNASeq results. *Cbx7* was among the differentiation genes that had increased transcripts at P14 compared to P7. Levels of Cbx7 proteins increase in the inner EGL and further in the IGL (Fig S1E), where GNPs have recently exited the cell cycle and initiated the first stages of differentiation.

GNP division, differentiation, and migration can occur during a wide developmental window, from P3 to P10 (*19*). The densely packed nuclei within the EGL and IGL preclude accurate quantification of protein levels. To avoid these problems we used GNP culture, which permits accurate and automated single-cell quantification. The murine *Math1* bHLH transcription factor was used to distinguish dividing from non-dividing cultured GNPs. *Math1* expression stops when the GNPs stop dividing. *Math-1* reporter GFP (*30*) fluorescence allows classification of GNPs as differentiated or proliferating. To estimate the changes in Cbx7 during differentiation in culture we compared 6h Atoh1 high cells (dividing GNPs) to 48 h Atoh1 low cells (differentiated GNs). Cbx7 was 3.3 times higher in the differentiated GNs then the dividing GNPs (Fig S1 D-G).

The transcription of the 787 differentiation genes varies in the exact timing of transcription activation, as reflected in Fig S1A. A regulator of a set of differentiation genes, among the 787, might be a good target for therapeutic manipulation to force MB cells to differentiate. Based upon gene expression patterns, SHH-subgroup MB cells are highly similar to Shh-exposed dividing P7 GNPs (*31*). Both cell types rely on Hh signaling as a mitogen (*1*). Mutations of Hh components in the GNP lineage are adequate to induce medulloblastomas (*6*). Further experiments examining epigenetic regulators therefore focused on P7 GNPs.

### The chromatin repression modification H3K27me3 is present in P7 GNPs on half the 787 differentiation genes, and correlates with H3K4me3 and poised RNA Pol2

To gain insight into the transcriptional regulation of differentiation genes, we investigated the chromatin state of P7 GNPs. The positions and abundance of six histone modifications and two chromatin-associated proteins were measured using ChIP-Seq: H3K27me3, H3K27ac, H3K36me3, H3K4me3, H3K4me1, H2Aubi119, Ring1b, and Pol2S5. We selected the histone modifications and known chromatin bound proteins for ChIP based on known markers of promoters, enhancers, and to broadly assess transcriptional repression.

Modifications associated with Polycomb-mediated transcriptional repression were mapped, including H3K27me3, which is produced by the PRC2 complex (*34*), H2Aubi119, which is produced by the PRC1 complex (*35*), and Ring1b, which is a core component of the PRC1 complex and is the ubiquitin ligase that catalyzes H2Aubi119 modifications.

Another form of transcriptional repression employs CpG island DNA methylation. DNA methylation was measured using methylated-CpG island recovery assay (MIRA), which uses recombinant MBD protein to bind double-stranded methylated DNA (*36*).

Protein modifications at active promoter regions were assessed, including H3K4me3, S5-phosphorylated RNA polymerase 2 (Pol2S5), (which labels paused RNA polymerase), H3K4me1 (which is associated with active enhancer regions), and H3K27ac (which is found at active promoters and enhancers) (*37–40*). DNA accessibility was assessed by ATAC-Seq, which is based upon the bias for Tn5 transposition towards open chromatin (*41*). To quantify the amount of a histone modification at a given gene we quantified the number of reads within 250 BP of the transcriptional start site (TSS). Several of the histone modifications, including the transcriptionally repressive modifications accumulate at the TSS and extend towards the gene body and upstream (Fig S2C,D). This extension of the histone modifications has been shown to be biologically relevant for H3K27ac at super-enhancers (*42*) and for H3K27me3 at permanently repressed genes like Hox genes. To better separate strongly marked genes from moderately marked genes we extended the quantified region to include significantly bound DNA regions farther from the TSS (Materials & Methods).

Of all the histone modifications and histone-associated proteins assessed for selective presence with the differentiation genes, the frequent presence of H3K27me3 was most striking. Of the 787 differentiation genes 447 (57%) had H3K27me3 within 250 BP of the TSS (Fig 2B). To identify important modifications we calculated enrichment, meaning the amount of a modification in the 787 differentiation genes compared to the amount in either all genes or in expressed genes during the GNP time-course. H3K27me3 was 2.3 times more frequent near differentiation genes compared to all expressed genes at P7 (p-val 7.54×10^-147^, hypergeometric BH adjusted) and 2.7 times more frequent compared to all genes (p-val 7.14×10^-198^, hypergeometric BH adjusted) (Fig. 2A). H2Aubi119, a repressive modification made by Ring1b, a component of the PRC1 complex, was also more frequent at the differentiation gene set with 2.17 enrichment over all genes (p-val 1.15×10^-245^, hypergeometric BH adjusted) and 1.89 over all expressed genes at P7 (p-val 3.71×10^-156^, hypergeometric BH adjusted).

**Figure 2:**
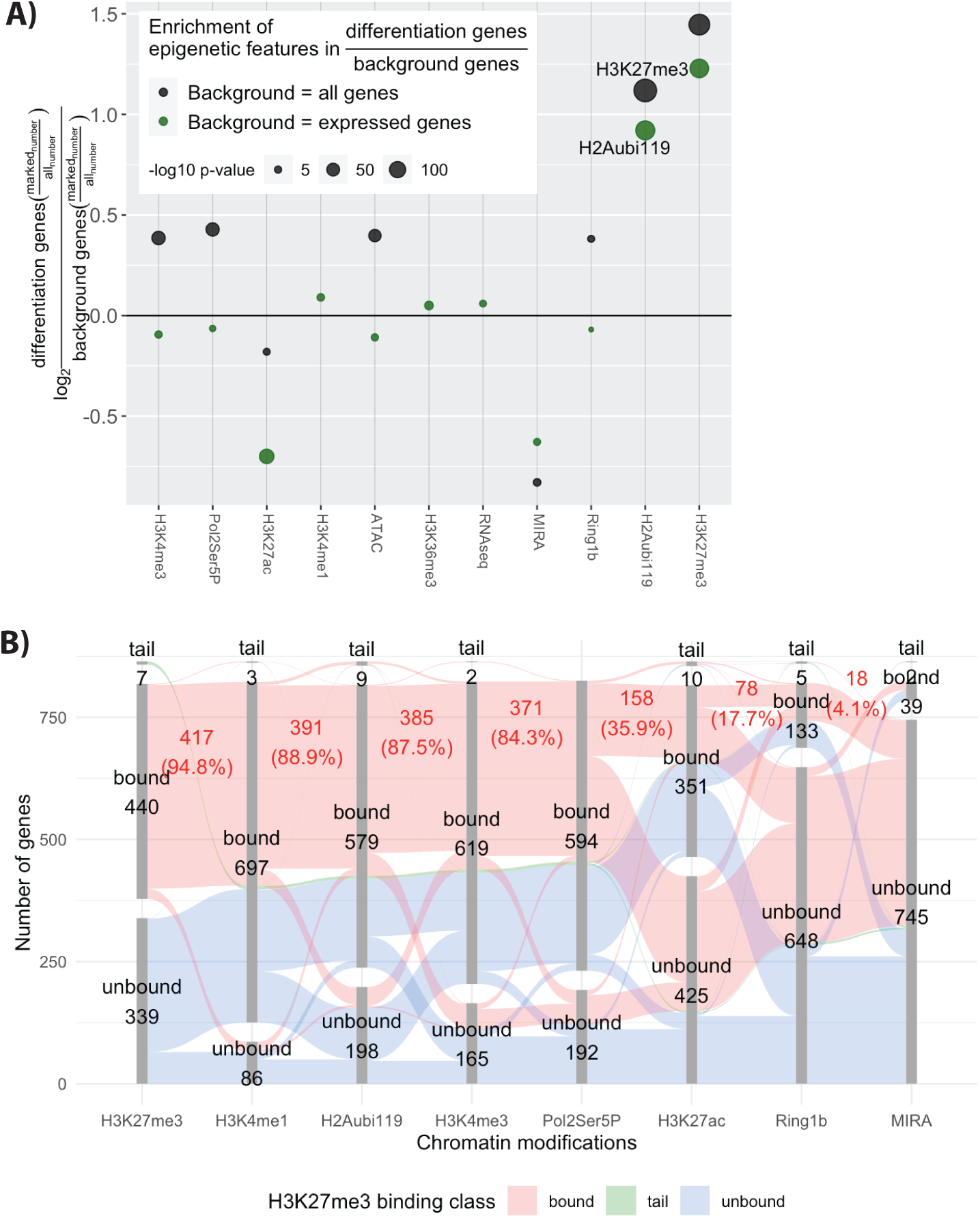
H3K27me3 modification near the promoter of genes at P7 is associated with genes that increase later with differentiation. A) Enrichment of chromatin features near promoters of proliferating P7 GNP whose expression increases during differentiation. Enrichment is calculated for a given modification as the number of differentiation genes over either all genes (black) or genes expressed at any point during the granule neuron lineage (green). The log2 value of this fraction is then shown on the figure. The p-value is calculated using the hypergeometric test and the p-values are shown as the size of the dot. **B)** A parallel sets diagram that shows correlation between H3K27me3 binding and other modification. The disconnected bar graphs shows the number of genes that are bound or within the long upper tail for each modification. The red colour that spans the bar segments shows the genes that contain the next modification.

The frequent presence of H3K27me3 modifications suggests that PRC2 may repress differentiation genes during GNP transit amplification. The PRC2 complex contains the methyltransferase EZH2, which carries out H3K27 methylation. From past studies (*43*) we know that PRC2 is present at specific chromatin regions, where it functions as a transcriptional repressor. The amount of H3K27me3 present regulates whether repression is temporary (moderate levels) or permanent (high levels) (*44–46*). The differentiation genes have a moderate amount of H3K27me3 (Fig S2A-C). Only three differentiation genes had high levels of H3K27me3. The other, non-differentiation, genes within the high H3K27me3 group included Hox cluster and transcription factors associated with alternative fates or earlier lineages. As an example, *Cbx7*, a differentiation gene, has moderate K27me3 while Lhx9 and Hoxa10 have high K27me3 (Fig SC). Very highly H3K27me3-modified genes are thought to be permanently repressed (*47*).

In addition to the amount of H3K27me3, whether PRC2 repression is transient or long-lived depends on its association with other histone marks or protein complexes. For example, H3K27 methylation may occur in association with active-gene modifications such as H3K4me4 and/or poised RNA polymerase II (*48*). This kind of arrangement is known as a bivalent promoter, transcriptionally repressed but poised for increased transcription (*49, 50*). In contrast, PRC2 can establish more permanent repression when associated with certain PRC1 variants (*51*). However, these are heterogeneous complexes, and changing select PRC1 subunits allows the association with bivalent promoters as well (*52*). H3K27me3 can recruit the machinery for DNA methylation (*53*), driving permanent repression.

To distinguish possible roles of PRC2 in GNPs we examined the relationship between H3K27me3 and H2Aubi119 (a marker of PRC1 complex) or H3K4me4 (a marker of active promoters) at differentiation genes that have H3K27me3. H3K27me3-marked differentiation genes have both H3K4me3 and PRC1 complex (H2Aubi119). DNA methylation was detected in only 4 percent of differentiation genes, indicating that this form of repression is mostly not active there.

The association of H3K27me3 with other modifications at differentiation genes is represented as a chart (Fig 2B). Each modification is represented by a column that shows the numbers of modified and unmodified genes. The columns are organized from left to right according to the number of genes that are also marked by H3K27me3. The red band shows genes containing H3K27me3 which share other modifications. Of the 447 H3K27me3-marked differentiation genes, 87 % also had H3K4me3 and 84.3% had paused RNA polymerase. PRC1 occupancy on the H3K27me3-bound differentiation genes was frequent: 89 % of the 447 genes had significant H2Aubi119. The frequent associations between H3K27me3 and H2Aubi119 or H3K4me3 are also seen with a scatter plot (Fig S2 D,E), where 373 (83%) of the differentiation genes bear all three modifications.

Thus at P7 many of the 447 differentiation genes had nearby PRC2 complexes, PRC1 complexes, and markers of active promotors such as H3K4me3 and RNA Pol2Ser5P (paused RNA polymerase 2). These results suggest that about half of the differentiation genes have bivalent or poised promoters containing both PRC2 and PRC1. Removal of these modifications in the normal course of development, or engineered removal in MB cells, may cause de-repression of these genes and early differentiation.

### Ezh2-mediated H3K27 methylation delays differentiation of GNPs

The H3K27me3 histone modification, associated with gene repression, is enzymatically carried out by the protein complex PRC2. The presence of H3K27me3 modifications at 447 GN differentiation genes at the P7 stage suggests that PRC2 may be blocking differentiation. In that case differentiation of GNs, or perhaps MBs, would require relief from PRC2 repression. We tested this hypothesis using mice carrying a mutation in *Ezh2*, which is the enzymatic component of PRC2 that catalyzes formation of H3K27me3. Homozygous *Ezh2* knockout mice die as embryos (*54*), so studying postnatal cerebellar development required conditional knockout mice. Mice with LoxP sites flanking the DNA encoding the SET domain of *Ezh2* (*55*) were crossed with *Math1-Cre* mice (*56*). The SET domain is required for Ezh2 to function as a methyltransferase (*57*) and maternal depletion of the SET domain in mice causes significant growth retardation (*58*).

As a consequence of crossing the *Ezh2* Lox-P mice with the *Math1-Cre* mice, H3K27me3 is reduced in the granule neurons (Fig S3 A-C). *Ezh1*, a paralog of *Ezh2* that is present throughout GN development is a weaker methyltransferase than *Ezh2* (*59*). Even with *Ezh1* functioning, conditional loss of *Ezh2* led to dramatic loss of H3K27me3 from GNPs and GNs, as shown in P7 cerebellar slices (Fig S3 A panel II vs VI, yellow arrow) and cultured GNPs (Fig S3 C panel II vs IV, yellow arrow).

The most notable phenotype of the *Ezh2* cKO mice is a blurring of the boundary between the inner external granular layer (iEGL) and the molecular layer (ML; Fig 3B-C, magenta and cyan arrow). GNs within the inner EGL (iEGL) permanently exit the cell cycle and remain there as they extend parallel fibers. GNs within the iEGL express the cell cycle inhibitor p27. In the *Ezh2* cKO the iEGL is more diffuse and blends into the ML (Fig 3A compare cyan arrows). To facilitate quantification of the EGL blurring we collapsed 2D fluorescent images into line segments (Fig 3D). P7 cerebellar slices were stained with anti-p27 to label post-mitotic EGL cells, NeuN antibody to identify the IGL, and DAPI to mark all nuclei. For each channel, measurements were made along a line from the pial border on the outside to the IGL. The 581 lines (266 WT and 315 *Ezh2* cKO) were aligned and averaged, in sets according to genotype (Fig 3E). The normalized p27 intensities show a spreading of iEGL, blurring the boundaries with the IGL and ML. The ML blurring, was quantified as an increase in the p27 at the junction of the iEGL and ML in the *Ezh2* cKO (Fig 3D; compare the cyan arrows). To quantify across replicates, we compared *Ezh2* cKO and WT at a point within the ML where the *Ezh2* cKO had the highest relative p27 fluorescence (Fig 3E).

**Figure 3:**
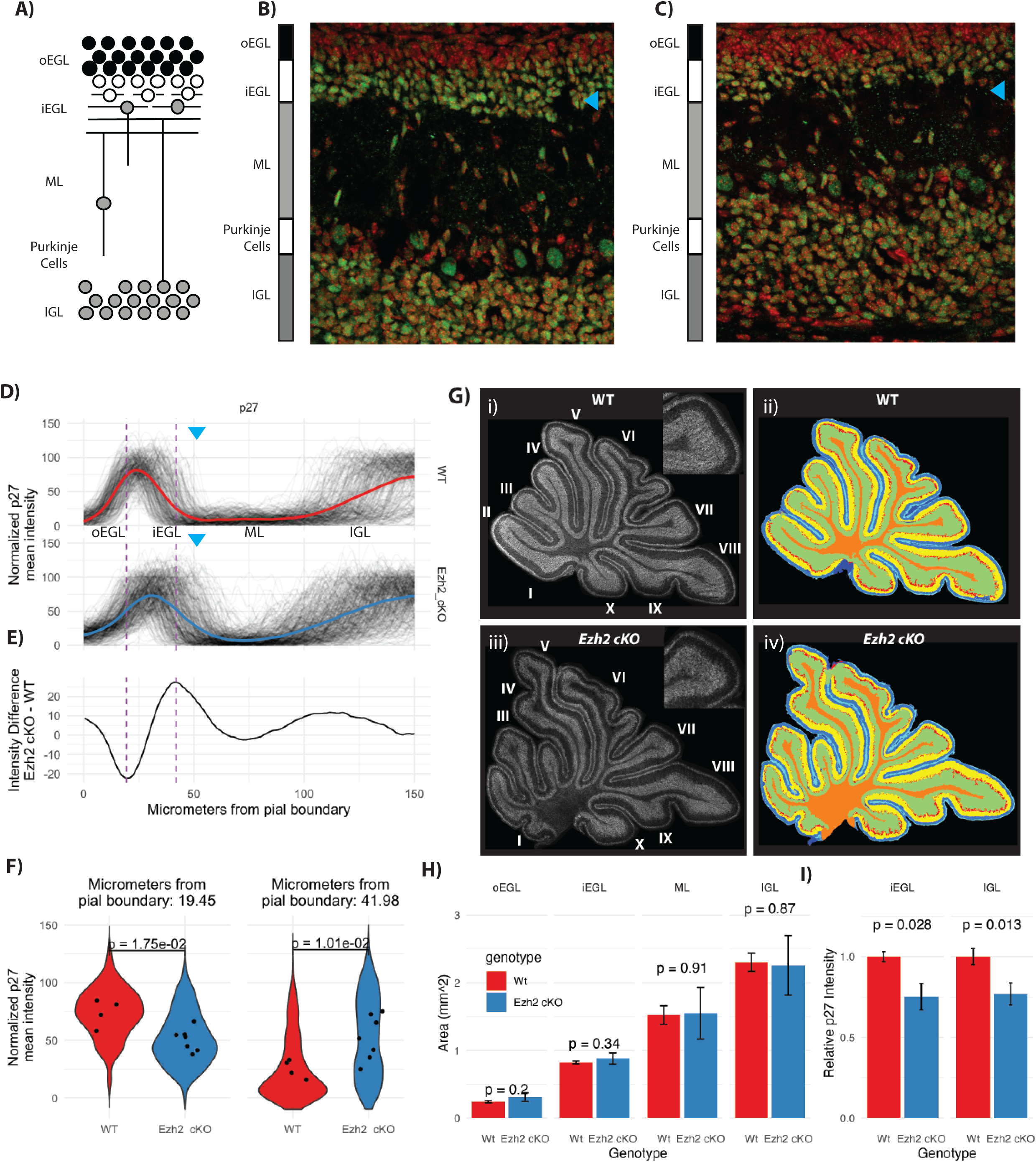
*Ezh2* cKO leads to a depletion of the inner EGL. **A)** Schematic of granule neuron (GN) development in a P7 cerebellum sagittal slice. Inner external grandule layer (iEGL), outer external granule layer (oEGL); molecular layer (ML), internal granule layer (IGL). **B)** WT and **C)** *Ezh2* cKO 40x confocal micrographs of the deep sulcus between lobule V and VI in the control and in the cKO, showing DAPI (red) and p27 (green; a cell cycle arrest marker in GNPs). In the mutant, the boundary between the inner EGL and the ML and the ML with the IGL is ill-defined. The cyan arrow indicates the position of the inner EGL, there are fewer p27 labeled cells sitting within the EGL. The cellular density is reduced in both the inner EGL and IGL. **D)** Quantification of the boundaries adjacent to the ML in WT and *Ezh2* cKO by collapsing 2D images into 1D line segments. The line segments for anti-p27 immunofluorescence span from the pial boundary to the IGL and show a hump of high average fluorescence at the inner EGL and the IGL. In the *Ezh2* cKO the iEGL p27 signal extends further into the ML then in the WT. The blue arrows show the approximate position of the transition between the iEGL and the ML. E) The average fluorescence intensity difference between mutant (n = 7) and WT (n = 4) shows where along the line segment from the pia to the IGL is the *Ezh2* cKO most different from the WT. Positive values are associated with p27 value being higher in the *Ezh2* cKO and is highest at the boundary of the inner EGL and the ML and to a lesser extent at the boundary between the ML and IGL. The WT shows higher signal within the inner EGL itself where the values of E are negative. **F)** Quantification of each line segment at the point where the differences between *Ezh2* cKO and WT are largest as indicated by the hashed lines seen in panel D and E. The left panel shows quantification at position where the WT had had more P27 staining then to the *Ezh2* cKO and the right panel shows the opposite. The distribution of the values at the 2 positions in right and left panel for the 266 WT and 315 *Ezh2* cKO line segments was shown as a violin plot (a histogram that is mirrored). The dots correspond to individual mice, with multiple slides analyzed for each mouse. P-values are calculated using a T-test showing a significant increase of P27 labeling within the boundary of the EGL and ML. G) Segmentation into inner and outer EGL, ML, IGL and deep white matter using panoramic images of the entire cerebellum, labeled with DAPI, P27, NeuN. Anti-p27 over the entire P7 cerebellar section, for WT i) and *Ezh2* cKO ii). **H)** Quantification of the area for each layer in WT and *Ezh2* cKO shows no significant difference, indicating the overall structure of the cerebellum is not dramatically altered and the phenotype is limited to the early migration out of the inner EGL. **I)** Quantification of the anti-p27 fluorescence intensity by layer shows a decrease in the fluorescence in *Ezh2* cKO versus WT, for the inner EGL (p-value 0.028, t-test) and IGL (p-value 0.013, t-test). The decrease in p27 abundance suggests that the total number of neurons within the inner EGL and the IGL density of cells within those areas is reduced.

The sizes of the layers of the cerebellum are largely unaffected by *Ezh2* cKO (Fig 3G,H). **T**he different layers of the cerebellum were measured in slices stained with anti-p27, NeuN antibody, and DAPI (Fig. 3F). No significant differences were observed in the thicknesses of the oEGL, iEGL, ML, or IGL (Fig 3G,H). Due to the diffuse EGL/IGL-ML boundary in the cKO, p27 fluorescence intensities within the iEGL (where GNPs begin differentiation) and IGL (where post-mitotic GNs reside) were reduced 25% and 23% (Fig 3I). These findings are consistent with the loss of *Ezh2* function not altering the overall structure of the cerebellum, but instead specifically affecting GN differentiation.

*Ezh2 cKO^-^* GNPs labeled with a 48 h EDU pulse migrated from the EGL prematurely (Fig. S3 D-F). To distinguish early migration out of the oEGL from stalled GN migration within the ML, GNPs going through S-phase were labeled with EDU and harvested 48 hours later. In WT mice there is dense EDU labeling of the iEGL, and GNs lining up along the border with the ML (Fig S3 D, panel III versus VIII, yellow arrows). In the *Ezh2* cKO, EDU**-**labeled cells are within the diffuse iEGL and within the ML (p-val 0.024), mirroring what was seen with p27 immunofluorescence (Fig S3E,F). We conclude that, in *Ezh2* mutant mice, GNs prematurely exit from the EGL into the ML, a movement typical of wild-type GNs that have stopped dividing.

The hypothesis that PRC2 blocks cell differentiation is consistent with the *Ezh2* mutant results, because early de-repression of differentiation genes would cause premature EGL cell migration.

GNPs cultured from *Ezh2* cKO mice, compared to normal, had more differentiated cells but the same fraction of dividing cells. That fits with the idea that loss of *Ezh2* allows premature activation of differentiation genes. To further test the impact of *Ezh2* cKO on differentiation, we cultured GNPs with Shh and monitored *Map2* (Fig. 4A,C), a marker for process extension, and *NeuN*, a marker of differentiated GNs. Process extension was measured using Neuroncyto 2 (*60*), which reveals cell processes and assigns them to a cell body (Fig. 4B,D). *Ezh2* cKO GNPs had a 37.6% increase (p-val 0.007) in average process length compared to control (Fig 4E).

**Figure 4:**
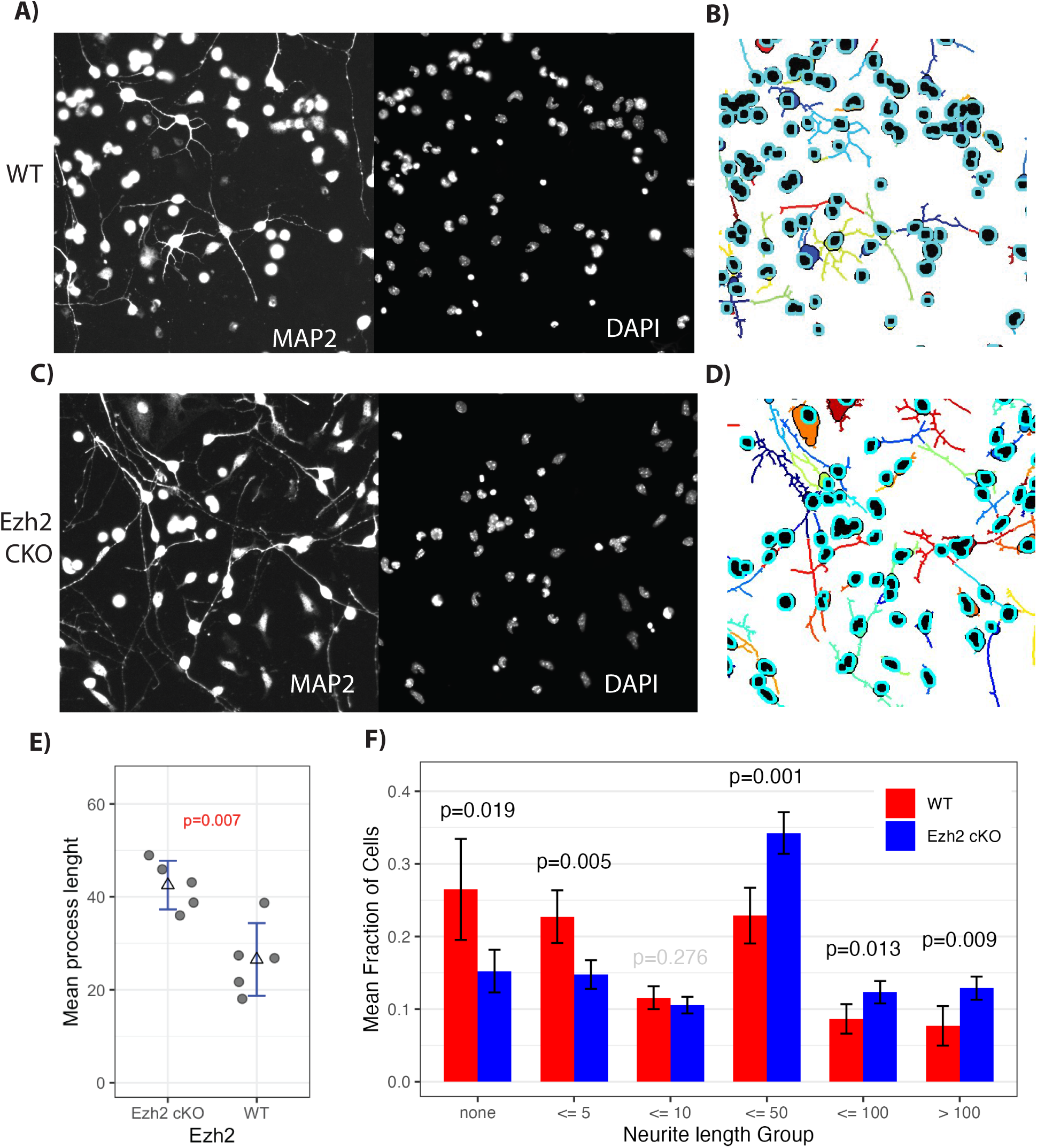
*Ezh2* cKO leads to premature process extension and NeuN expression. GNPs cultured for 24 in Shh, were labeled with DAPI (nucleus) and for MAP2 (neuronal processes) (**A,C**), then segmented with Neuronctyo 2 (**B,D**). Segmented images showed increased process extension in *<u>Ezh2</u>* cKO (**D** for cKO versus WT in **B**). **F**) Mean process length comparison between *Ezh2* cKO and WT shows cells in *Ezh2* cKO mice had significantly longer processes (n = 5). **E**) Quantification of the number of cells with no process, a process less than 5, 10, 50, 100 pixels, or greater then 100 pixels, showed a significant increase in the number of processes longer then 50 pixels. P-value determined by T-test.

Next, we separated the cells into groups based upon process length (Fig 4F). WT cells had a 19.2 % higher proportion of cells with no process (49.2 % WT vs. 30% in *Ezh2* cKO; p-val 0.019) or a process less than 5 pixels long (p-val 0.0047). *Ezh2* cKO cells had processes 10 pixels and longer 20.1% more often (39.3 WT versus 59.4 *Ezh2* cKO). Thus loss of *Ezh2* led to more of the longer processes, an indicator of differentiation, in cultured GNPs.

Cell cycle stage and differentiation were quantified using staining for DAPI (to show all nuclei), Ki67 (to show dividing cells), and NeuN (to show post-mitotic neurons) (Fig S4A). The fraction of total cells that were NeuN-positive in *Ezh2* cKO cultures was 13.9 % compared to 9.8 % in WT (p-val 0.008) (Fig S4C). The amount of NeuN fluorescence increases on the per-cell histogram (Fig S4B). No significant change in the number of cells that were in G1 (p-val 0.107) or G2 (p-val 0.087) occurred, with cells more often being in G1 in *Ezh2* cKO and in G2 in WT (Fig. S4C). *Ezh2* appears to be a cell-intrinsic regulator of differentiation timing; its loss hastens process extension and nuclear migration out of the EGL.

### During normal development, *Ezh2* transcript and protein levels decrease as GNPs differentiate

Reduced PRC2 activity as GN differentiation commences could be due either to reduced Ezh2 protein levels or to functional inactivation. Among genes encoding components of repressive chromatin complexes, transcript levels for *Ezh2* and *Cbx7* changed the most during early GN development, with *Ezh2* transcript being reduced 8.8 fold after P7 (Fig 5A,B). GNP cell cycle exit coincides with a reduction in Ezh2 protein level, as shown by reduced staining intensity in the iEGL (Fig 5C). To further explore this observation, P7 GNPs freshly isolated from *Math1>GFP* reporter mice were plated to observe differentiation. As early as 6 hours after plating, the average level of Ezh2 was lower in GNs that had begun to differentiate, as determined by low Math1-GFP levels. Relative fluorescence for Ezh2 continued to drop at 24 hours of culture (Fig 5D). Thus cell cycle exit strongly correlates with decreased Ezh2 levels in cultured cells and in vivo.

**Figure 5:**
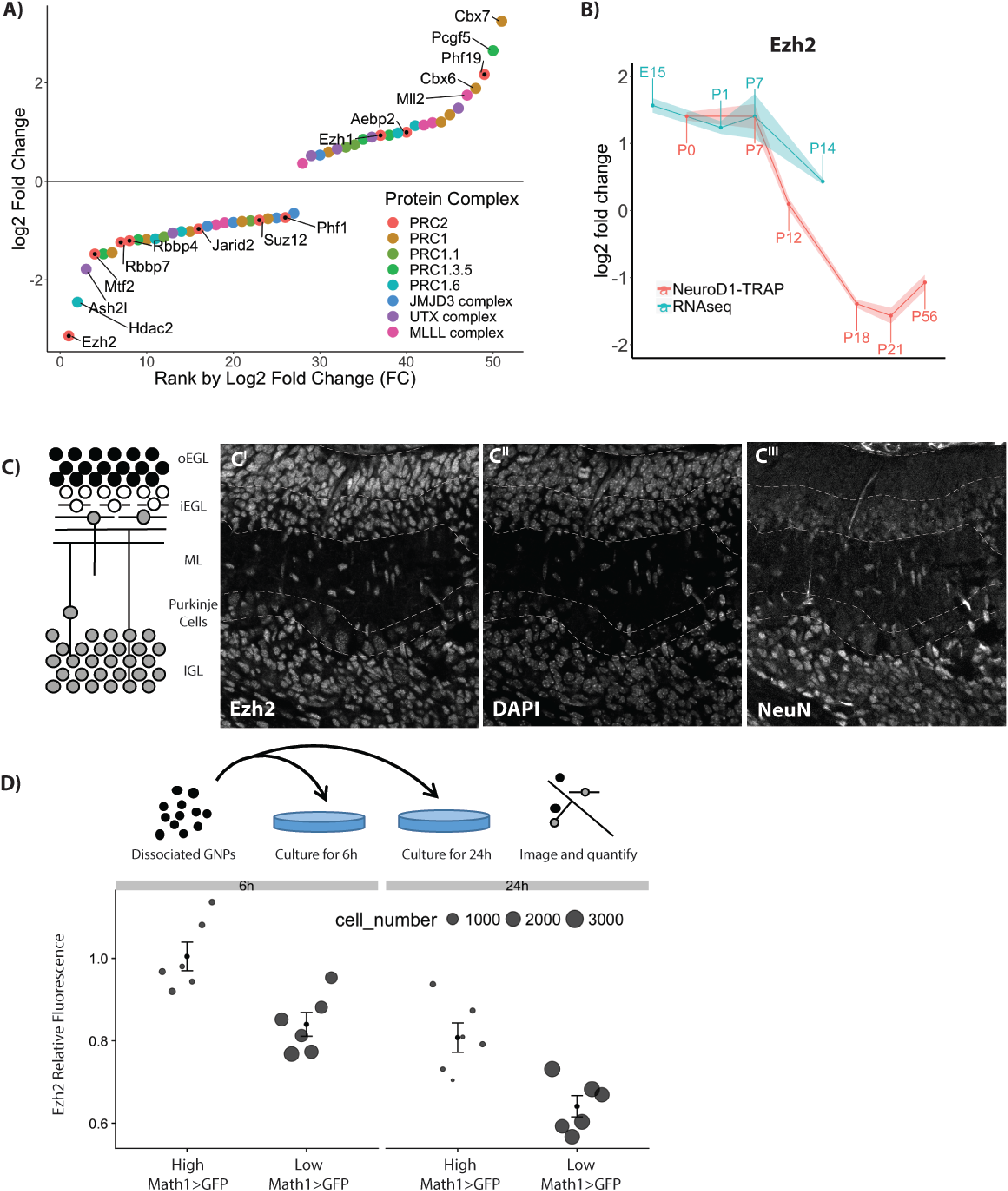
Transcriptional regulation of known H3K27me3 effector protein complexes during granule neuron (GN) development. **A)** Rank order plot of genes that change expression during GN development shows *Ezh2* as the gene with the largest decrease in RNA abundance. The y-axis indicates the combined fold change for both datasets. **B)** RNA abundance over time for *Ezh2*. **C**) Schematic of granule neuron (GN) development in a P7 cerebellum sagittal slice. A P7 cerebellar slice stained with Anti-Ezh2 (left), DAPI (middle), and NeuN (right) shows reduced Ezh2 staining intensity in the postmitotic cells of the iEGL and EGL. **D)** Schematic of GNP culture without Shh, which causes GNPs to immediately exit cell cycle. Ezh2 protein abundance was measured after 6 h and 24, splitting cells into proliferating (high Math1>GFP) cells and non-proliferating (low Math1>GFP) cells (n=6). The decrease in Ezh2 shortly after cell cycle suggests that the decrease in Ezh2 is linked to cell cycle exit but does not show a causal relationship.

The results so far suggest that Ezh2-based H3K27me3 modifications at promoter regions delay transcription of many differentiation genes during GNP proliferation. Once GNPs complete their final mitotic division and enter G0 in preparation for differentiation, reduced *Ezh2* function may foster activation of the GN differentiation program. Can this knowledge be applied to manipulate MB cells?

### Medulloblastoma cells also H3K27-trimethylate GN differentiation genes

In MB cells, as in dividing GNPs, repressive chromatin regulators may prevent expression of GN differentiation genes to maintain a pro-proliferative state. Indeed, 95% of the 447 differentiation genes that are H3K27-marked in GNPs are repressed in MB cells compared to GNs (Fig 6C). To determine whether GN differentiation genes are stably repressed in MB, we measured chromatin markers and mRNA transcript abundance in MB samples from *Ptch1^+/-^* mice. *Ptch1* encodes the Hedgehog receptor, a negative regulator in the pathway, so in these mice, and in humans with the same kind of mutation, derepressed Hedgehog target genes increase the frequency of MB. RNAseq was used to compare transcript populations in MBs from *Ptch1^+/-^* mice to P1, P7, and P14 GNPs. As expected, MB transcripts correlated most with transcripts from rapidly dividing P7 GNPs (R^2^ = 0.79) compared to P1 (R^2^ = 073, Fig S6 B) or P14 (R^2^ = 0.52, Fig S6 C).

**Figure 6:**
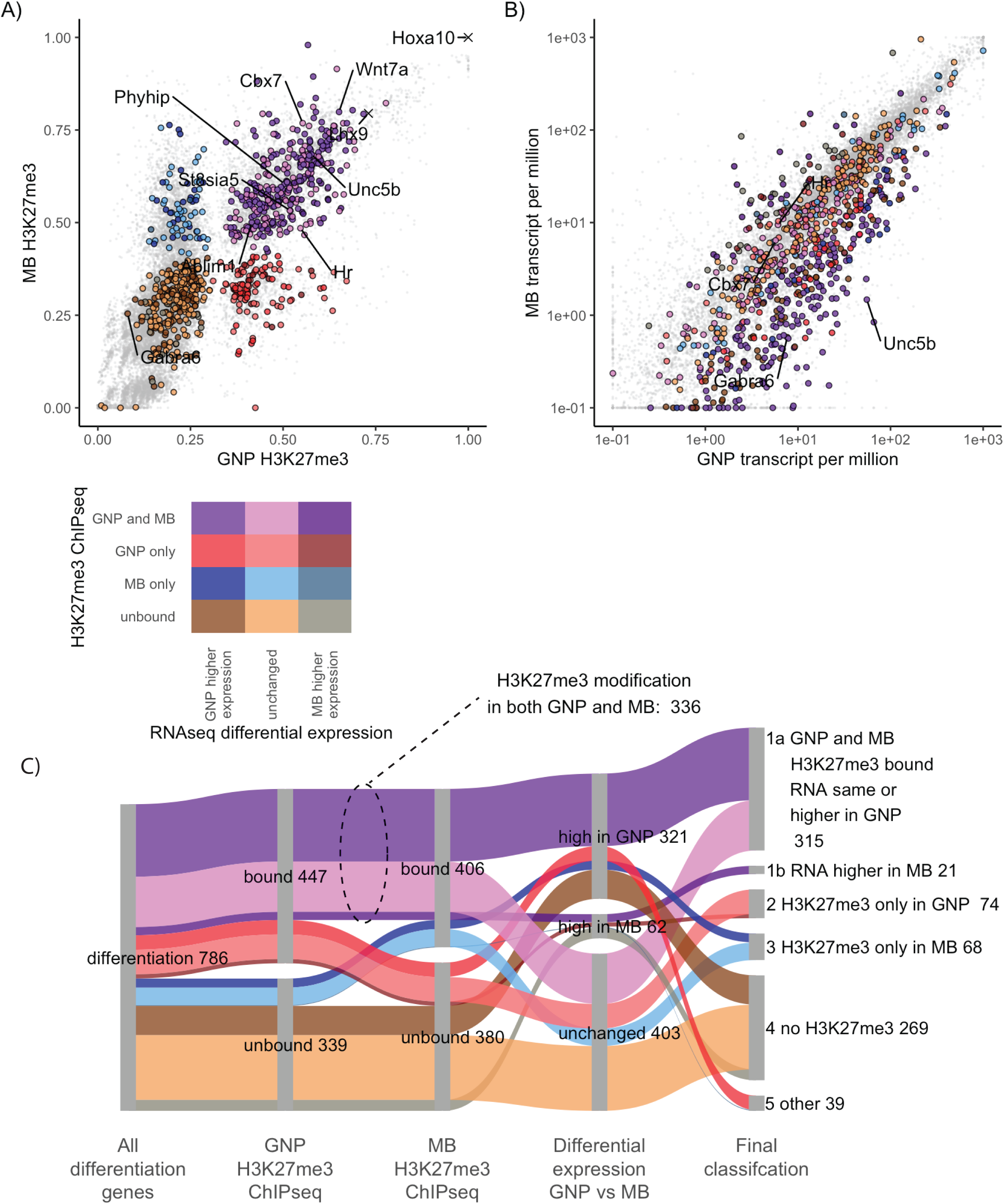
In medulloblastoma cells GNP differentiation genes are transcriptionally repressed and H3K27me3 modified. **A)** H3K27me3 quantified over the promoters of GNP differentiation genes in GNPs (x-axis) and MB (y-axis). Genes are color-coded based upon H3K27me3-marked versus not marked in GNP and MB. Genes whose promoters show more H3K27me3 modification in GNP are shown in red, while those higher in MB are blue and those modified in both are purple. Duller colors (dark tones) were used if higher relative expression in MB than in GNPs. Higher saturation colors indicate gene expressed relatively higher in GNPs than in MB. The colour legend is seen bellow panel A. **B)** Transcript levels of differentiation genes in P7 GNPs, x-axis) versus in *Ptch1*^+/-^ derived MB cells (y-axis). Colors are inherited from panel A with dark tones used for genes in MB (2 fold higher expression and adjusted p-value <0.05) and high saturation colours when higher expression in GNPs. **C)** Sankey diagram showing how the H3K27me3 modified differentiation genes are expressed in MB. The second column shows the differentiation genes in GNP separated by H3K27me3 modification. The third column compares MB H3K27me3 ChIP, showing of 447 H3K27me3 modified GNP differentiation genes, 336 (75%) of those genes are also H3K27me3 modified in MB. The fourth column shows differential expression between P7 GNPs and MB. Even when the H3K27me3 is not present in MB cells only 8% of the differentiation genes show an increase in expression in MB compared to P7 GNPs. The fifth column shows the final classification.

In MB cells, three-quarters of the GN differentiation genes have H3K27me3 at their promoters (Fig S6 A,C). The Pearson correlation between MB cells and P7 GNPs was also strong for histone markers, with H3K27me3 being 0.79, H2Aubi119 being 0.87 and H3K4me3 being 0.95 (Fig. S6A). Among genes that increase expression during GN differentiation, only 27% of the H3K27-methylated GN differentiation genes did not have H3K27me3 in MB cells (Fig 6 A,C). The majority of GN differentiation genes are persistently repressed in MB.

### In human SHH-subtype medulloblastomas, higher levels of *EZH2* transcript correlate with lower-level expression of GN differentiation genes

The inverse relationship between GN differentiation gene transcription and Ezh2 levels holds true in human MB tumors as it does in mouse MBs and developing mouse cerebellum. The levels of *EZH2* transcription from 223 primary human samples of *SHH*-subtype MB (*9*) were compared to the levels of GNP differentiation gene transcripts. Samples were ranked in terms of *EZH2* mRNA level (Fig 7A). The 30% of the tumors with the least *EZH2* RNA were compared to the top 30% (Fig S7A). GN differentiation gene transcript levels were on average 1.3 fold higher in tumors with low EZH2 transcript than in tumors with high EZH2 (Fig S7A).

**Figure 7:**
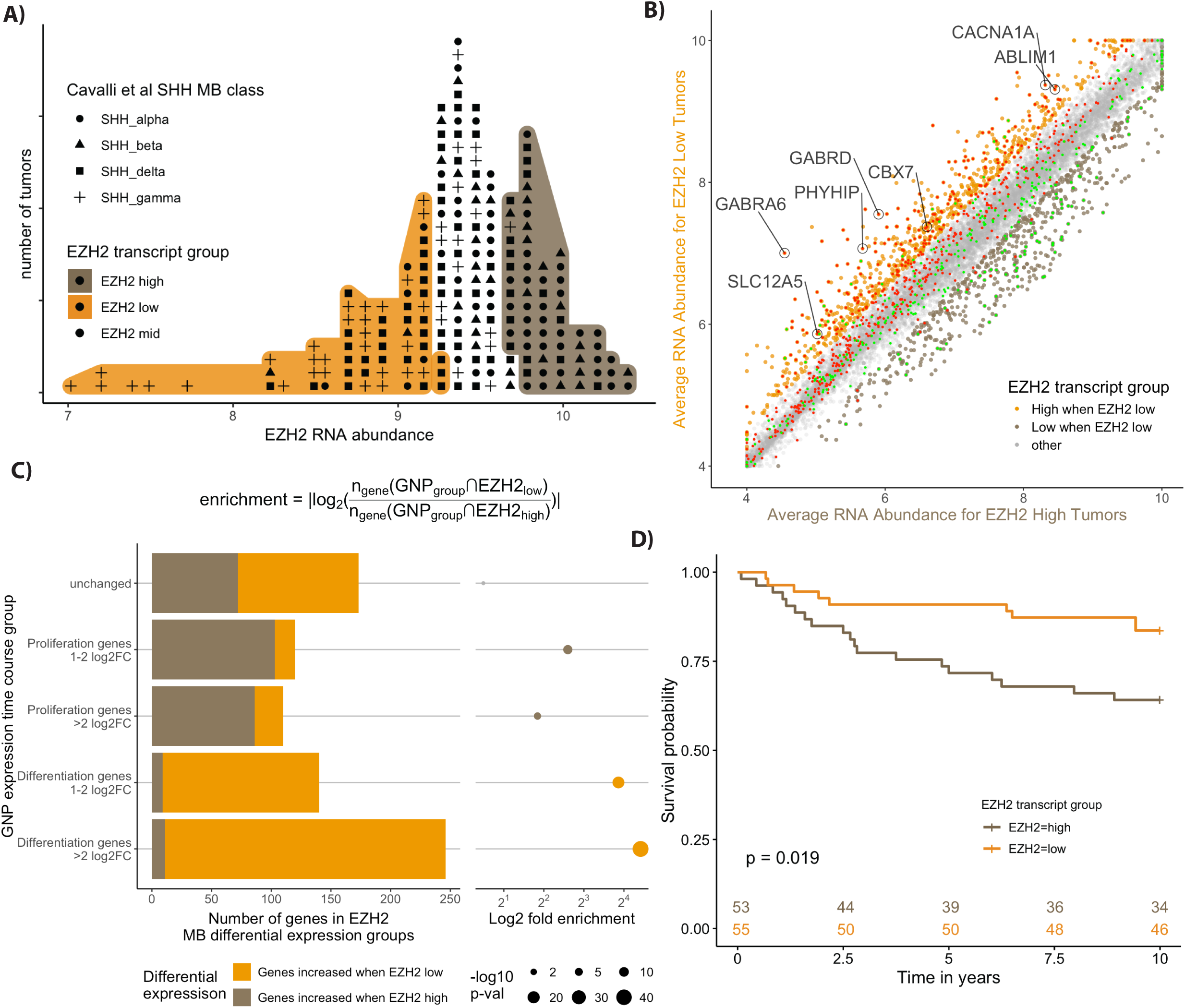
Human SHH MBs with High EZH2 are associated with worse outcome and show lower expression of GNP differentiation genes. **A)** Labeled dot histogram of *EZH2* transcript abundance from 223 human SHH MB with the top (orange) and bottom (brown) 30% labeled. Each human tumor is represented as a dot with the symbols reflecting the most recent SHH MB classification (*9*). Note the high number of patients with gamma-class SHH MB with low EZH2 expression. These patients are infants with a typically good prognosis. **B)** Comparison of average gene expression between the human MB samples from the 30% highest (x-axis) and lowest EZH2 (y-axis) expressing tumours. The color labels show more highly expressed genes that are 1.5 fold higher (adjusted p < 0.05) in tumours that were low in EZH2 were labelled orange, while those that are higher in *EZH2* high tumours are brown. The GNP differentiation genes (red) show considerable overlap with the genes highest in EZH2 low tumours (orange), while the GNP proliferation genes (green) show overlap with the genes that are highest in EZH2 high tumours (brown). **C)** Quantification of the extensive overlap between GNP differentiation genes and genes highly expressed in EZH2 low human MB tumours. The categories of genes based on the GNP time course (y-axis) are quantified using a bar graph that shows the intersection of genes for each GNP group to either genes that are highly expressed when Ezh2 is low (EZH2_low_) or genes that are highly expressed when Ezh2 is high (EZH2_high_). Enrichment is calculated as the number of GNP group genes that intersect (∩) with EZH2_low_ genes over the number that intersect with EZH2_high_ genes as described in the equation. The p-value is determined by the hypergeometric test. **D)** Kaplan-Meyer curves show a significant 10 year survival difference for SHH MB patients with high versus low EZH2 transcript.

Comparing human genes that were differentially expressed in *EZH2*-high vs *EZH2*-low SHH MB samples with the expression of their mouse homologs revealed a striking pattern (Fig 7B). Genes that were significantly more highly transcribed in human MBs that had relatively low *EZH2* RNA were 26-fold enriched for genes whose transcription increases during mouse GNP differentiation (Fig 7C). 60% of the human genes that had low expression when *EZH2* RNA was high were among the mouse GNP differentiation genes (343/570). The amount of Ezh2 transcript in each individual tumour was also anticorrelated with the transcript levels of the individual differentiation genes (Fig S7B,C). The average Pearson correlation for the differentiation genes was -0.24 indicating a negative correlation (Fig S7B,C). Thus the relationship between *EZH2* transcription and differentiation gene transcription is maintained in human MBs.

Patients with high *EZH2* RNA tumors had significantly worse 5-year (*EZH2*-high 73% versus *EZH2*-low 91%) and 10-year (*EZH2*-high 64% versus *EZH2*-low 84%) survival (Fig 7D), so patients with lower *EZH2* RNA and more differentiation gene expression have a better prognosis.

### Combined Ezh2 inhibitors and CDK4/6 inhibitors force MB cells to undergo neuronal differentiation

Mouse MB cells isolated from *Ptch1*^+/-^ mice (*61*) were plated on laminin-coated plates and cultured for 72 hours in serum-free medium, with or without small molecule inhibitors (Fig 8A). We measured rates of proliferation and differentiation using 2 combinations of antibodies and dyes. In one type of experiment, we used anti-pRb (Ser807/811) to label G1/2 versus G0 and NeuN to label differentiated cells (Fig 8A,B). In a second type of experiment, the antibody combination was p27, which identified quiescent G0-arrested MB cells, and Atoh1, which is only present during MB proliferation (Fig S8A,B). Cycling MB cells have high Atoh1 and low p27 if in G1/S/G2 (green in Fig. S8A) or high p27 if in G0 (yellow in Figure S8A). Cells embarking on differentiation (red in Fig. S8A) have high p27 but relatively low levels of Atoh1. NeuN marks a later stage of development than p27 presence or loss of Atoh1.

**Figure 8:**
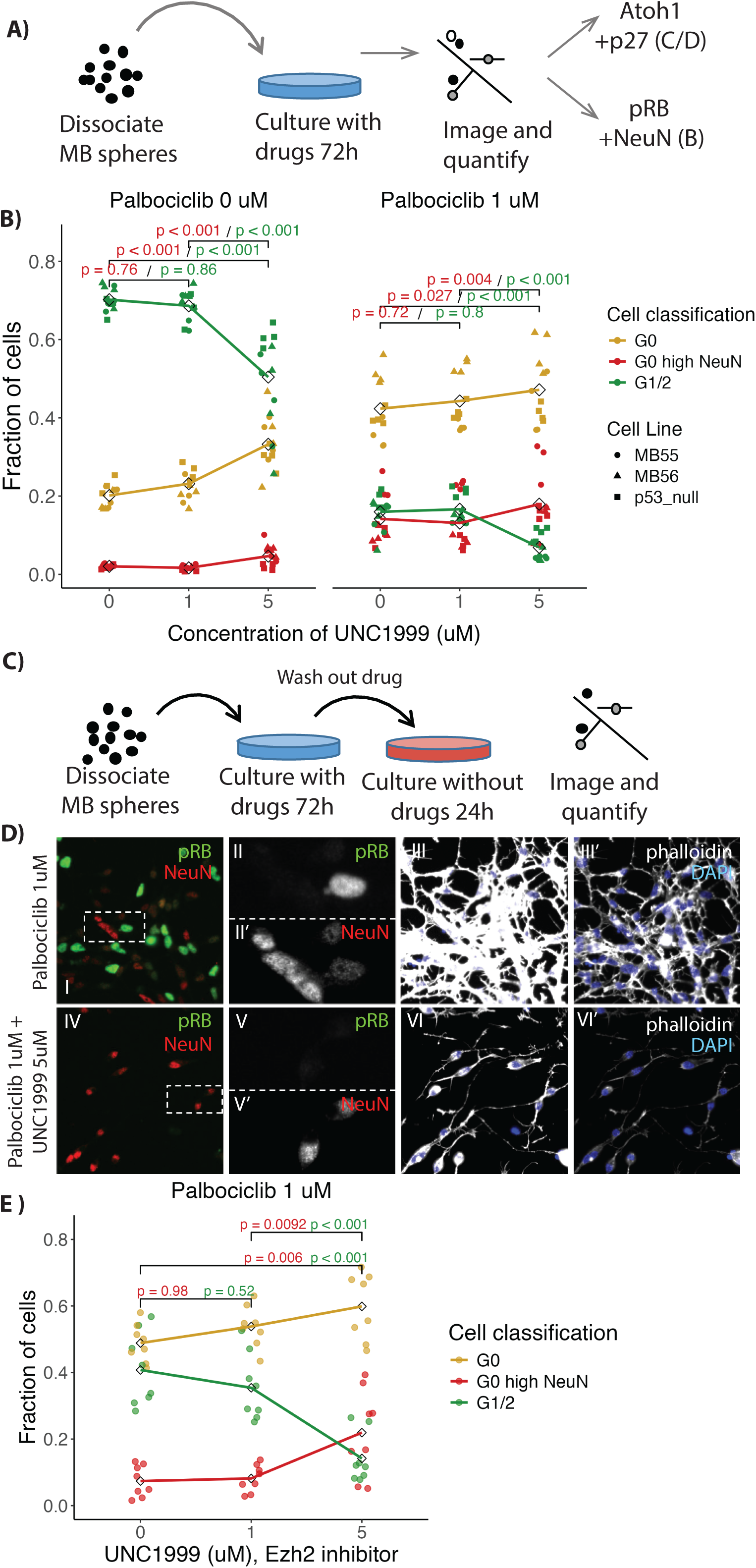
Treatment of MB cells with Ezh2 and CDK4/6 inhibitors leads to differentiation. **A)** Schematic of treatment and imaging protocol. MB cells were treated using the Ezh2 inhibitor UNC1999 with or without the CDK4/6 inhibitor palbociclib. **B)** Quantification of immunofluorescence in treated MB cells (n = 16), derived from *Ptch1*^+/-^ mice, stained for NeuN, pRb (Ser807/811) and DAPI to categorize the cells as in G1/2 (green), G0 (yellow) and differentiating (G0 with high NeuN (red)). Note an increase in fraction of cells scored as differentiating with combined Palbociclib and UNC1999 treatment. **C)** Schematic of treatments with inhibitors followed by drug wash out. **D)** Immunofluorescence images of fields of MB cells stained for NeuN (red), pRb (green), DAPI (blue) and phalloidin (grey) after treatment for 72h with 1 uM Palbociclib (i-iii) combined with 1 uM Palbociclib or 5 uM UNC1999 (iv-vi) followed by washout as per C. Dashed boxes in i and iv show regions enlarged in ii and v respectively, with the individual colour channels shown in separate panes. Note fewer pRB positive cells with UNC1999 and Palbociclib (iv) than with Palbociclib alone (i). **E)** Quantification of fractions of cells in G0, G1/2, and G0 and differentiating as described in B (n = 8). Note that the MB cells treated with both UNC1999 and Palbociclib showed persistent cell cycle exit even after washout. P-values in B and E calculated using ANOVA with post-hoc Tukey test.

Untreated MB cultures had 70.2/43.6% (pRb high / Atoh1 high p27 low) dividing cells and 2.0/14.2% (NeuN high / Atoh1 low p27 high) differentiating cells. Cultures treated with Ezh2 inhibitor (5 uM UNC1999) had 50.4/18.6% (pRB high /Atoh high p27 high) dividing cells and 4.6/ 35.2% (NeuN high / Atoh1 low p27 high) differentiating cells (Fig 8B, Fig S8B). Ezh2 inhibition caused a 21 % increase in the fraction of differentiated cells, based upon measuring Atoh1 and p27 (p-val < 0.001) and a 2.6 % increase in NeuN positive cells (p-val < 0.001). For UNC1999, previous studies of cultured cells showed that a dose of 1 to 5 uM caused substantial reduction in H3K27me3 activity without substantial toxicity (*62*). Thus induction of differentiation genes using Ezh2 inhibitor reduced but did not stop MB cell growth, in keeping with a lack of effect on MB frequency due to genetic removal of Ezh2 function. *Ptch1^+/-^* mice were crossed with *Ezh2^fl/fl^*Math1-Cre mice. About 15% of the *Ptch1^+/-^* mice develop MBs. *Ezh2* cKO did not affect the rate of MB formation in *Ptch1^+/-^* mice. MB formation occurred in 14.86% (n = 74) who were Cre^+/-^ and 14.28 % (n = 84) in Cre^-/-^ animals.

Normal development may be informative about how to stop MB cells from growing. During normal differentiation, GNPs would first enter G0 arrest, then GNP differentiation genes would be de-repressed as Ezh2 is reduced. To stop MB cell proliferation, the same two steps might be required: going into G0 arrest and then differentiating. MB cultures treated with CDK4/6 inhibitor (1 uM Palbociclib) to drive them into G0 (*63*), had 18.1/15.4% (pRb high / Atoh high p27 low) dividing cells and 15.9/26.0 % differentiating cells (NeuN high / Atoh1 low p27 high). Thus an inhibitor that pushes MB cells toward G0 is does increase rates differentiation more than the Ezh2 inhibitors alone but still at relatively low rates (Fig 8B, Fig S8B).

Dual treatment with Ezh2 inhibitor and CDK4/6 inhibitor produced the highest percentage of terminally differentiating MB cells 22.8/43.0 % (NeuN high / Atoh1 low p27 high) with only 9/6.8 % (pRb high / Atoh1 high p27 low) percent of cells dividing (Fig 8B, Fig S8B). The combination of Ezh2 inhibitors and CDK4/6 inhibitors increased the rate of differentiation by 7% for NeuN positive cells (p-val 0.004) and 17% for low of Atoh1 with high p27 (p-val < 0.001). MB cells treated with both 5uM UNC1999 and 1um Palbociclib developed thin neuron-like processes, that contain the F-actin stain phalloidin (Fig S8B xiii). In contrast, after treatment with CDK4/6 inhibitor alone, no such processes were observed and the cells appeared similar to dividing MB cells (Fig S8B x). It should be noted that differentiated GN require changes to the media for long term survival which could limit the time an MB cell could survive late into neuronal differentiation.

Combining a CDK4/6 inhibitor with an Ezh2 inhibitor could be toxic to dividing cells. The increase in percentage of differentiated cells would then reflect more robust survival of differentiated cells vs proliferating cells. If selective survival explained the results, then the absolute number of differentiated cells should be similar in untreated and doubly treated cells. Instead, combining 5uM UNC1999 and 1 uM Palbociclib produced 2.2 to 4.68 times more differentiated cells compared to control cells not treated with any drug (Fig S8C). This suggests that combining Ezh2 and CDK4/6 inhibitors drove conversion of dividing cells into differentiated cells, rather than causing selective death of dividing cells.

A key advantage of differentiation therapy over anti-proliferative approaches is that terminally differentiated neurons should not re-enter the cell cycle even after the treatment is terminated. MB cells in culture were treated with 1 uM of the CDK4/6 inhibitor alone or in the presence of various doses of Ezh2 inhibitor. After 72 hrs, all drugs were washed out and the cells cultured for an additional 24 h (Fig. 8C). As previously, cells were labeled using anti-pRb (Ser807/811) and NeuN (Fig. 8D). When CDK4/6 inhibitor was used alone, following drug wash out 40.8 % of MB cells re-entered the cell cycle (Fig 8E). Thus, with only CDK4/6 inhibition, MB cells can rapidly re-enter the cell cycle. Only 14.2 % of cells treated with both inhibitors re-entered G1/2. Again, we saw an increase fraction of differentiated from 14.5 % of cells to 21.9% (p-val 0.006) (Fig 8E) and in the total number of differentiated cells by 1.68 times (p-val 0.016) comparing dual Ezh2 and CDK4/6 inhibitors compared to CDK4/6 inhibitors alone (Fig S8D). The combination of Ezh2 inhibitors with CDK4/6 mediated cell cycle arrest achieves the most important therapeutic goal, to irreversibly prevent cell cycle re-entry.

## Discussion

The parallels between early cerebellar development and MB formation provide an opportunity to study key regulators of normal development in the context of tumor cells, with the goal of stopping tumor growth. During normal cerebellar development, GNPs proliferate under the mitogenic effect of the Sonic hedgehog pathway. Following a period of transit amplification, GNPs exit the cell cycle and terminally differentiate into granule neurons, the most abundant neuron in the brain (Fig 1). In MBs, the SHH pathway is aberrantly activated through mutations that inactivate inhibitory elements of the pathway (eg Patched or Sufu) or through amplification of activating elements (eg *Gli2* amplification). As a result, cells fail to exit the cell cycle and can ultimately give rise to a tumor. A large body of evidence describes the morphological (*64*), transcriptional (*65*), and post-translational modifications (*66*) similarities between transit-amplifying GNPs and medulloblastomas.

### Ezh2-mediated H3K27me3 broadly represses differentiation genes in GNPS and MBs

Here we undertook a genome-wide identification of the epigenetic regulators and transcriptional changes that occur during the initiation of proliferation, peak proliferation, and early stages of differentiation in GNPs. In agreement with other groups, we find that many genes whose transcription increases as GNPs differentiate have bivalent promoters, with H3K27me3 and H3K4me3 modifications (*67, 68*). Our study also shows PRC1 occupancy at the majority of the H3K27me3-marked differentiation genes (Fig 2). Most PRC1 complex variants and the PRC2 complexes are involved in transcriptional repression, which allows a transient repression of genes required during differentiation. The roles of PRC2 and PRC1 complexes during differentiation has been well described in many different cell types including other types of neurons (*69*), myocytes (*70–72*), cardiomyocytes (*73*) and T-cells (*74*).

### Bivalent H3K27me3 and H3K4me3 modifications delay transcription of specific gene sets during GNP cell cycle exit

For a cell with complex cytoarchitecture like a neuron, it is important to turn on the correct genes during differentiation and to have those genes turn on at the correct time. Ezh2 is transcriptionally activated by the pRb/E2f1 complex during S-phase expression(*75, 76*). In GNPs the bivalent H3K27me3 and H3K4me3 modifications mark a subset of genes that appear to be sensitive to the drop in Ezh2 function. The subsequent transcriptional activation of genes associated with neuronal differentiation, like ion channels, can in this way be reliably linked to cell cycle exit. The reduction of Ezh2 protein (Fig 5) that occurs as GNPs complete their cell division stage is likely to promote terminal cell cycle exit by allowing de-repression of differentiation genes.

What determines the exact timing of transcriptional activation for a given differentiation gene is likely multifactorial. Kdm6b, an enzyme that removes H3K27me3 methylations, significantly influences timing during GNP differentiation (*77*). The abundance of Kdm6b across the H3K27me3 marked differentiation genes may be non-uniform, activating some genes earlier than others. The PRC1 complex has numerous variants, which could fine tune the required delays in GNP differentiation gene expression timing. Another path of H3K27me3 elimination can be replicative dilution, where H3K27me3 modifications are depleted by ongoing cell division following Ezh2 activity reduction (*45, 46*). In GNPs, the drop in *Ezh2* expression happens as cells exit the cell cycle, so replicative dilution does not seem to be an important factor.

### Ezh2 loss accelerates differentiation but only after cell cycle exit

*Ezh2* cKO allowed early differentiation of GNPs but did not have an impact on transit-amplifying GNPs. During GNP cell division, Ezh2 is highly expressed. As Ezh2 levels drop following cell cycle exit, GNPs activate H3K27me3 modified genes. Premature removal of Ezh2 should cause early and/or prolonged activation of differentiation genes. Our *Ezh2* knockout did result in early differentiation but only after the period of transit amplification was complete (Fig 3). In cultured GNPs, *Ezh2* cKO did not reduce the number of G1/2 cells. If anything, there was an increase in the number of G1 cells (Fig 4). The phenotype of the *Ezh2* cKO is quite subtle; a previous study did not report a phenotype in the cerebellum in *Ezh2* cKO (*78*). We too found it difficult to observe and quantify the changes to the iEGL. We were alerted to the phenotype by automated analysis of GNP cultures from *Ezh2* cKO mice that revealed increased process extension pointing our attention to the iEGL.

### H3K27me3-marked differentiation genes have known GN differentiation phenotypes

Many H3K27me3-marked differentiation genes have known neuronal differentiation phenotypes in cerebellar GNs. One important example is the voltage-gated Ca^2+^ channel genes, which have been linked to GN migration. Of the six channel genes that increase transcription during differentiation, five are modified by H3K27me3. Loss of function of these genes impairs normal GN differentiation. Knockout of *Cacna1a* which codes for Cav2.1 showed clear migration defect in the EGL, with persistence of the EGL at P21 (*79*). Another potent regulator of Ca^2+^ that is critical to GN migration is the NMDA receptor (*80*), which is a ligand-gated glutamate channel. The genes for all four NMDA subunits that transcriptionally increase (NR2BA-D), are modified by H3K27me3. Functional NMDA receptors form when the radially migrating GNs start migrating into the ML (*81, 82*). NMDA receptors undergo subunit switching between migrating and post migratory receptors (*83, 84*). Genetically altering subunit composition leads to an increased rate in migration (*85*) and persistent EGL (*86*), while blocking NMDA receptors reduces migration(*87*). The targets of Ezh2 have developmental phenotypes, and the timing of their production matters.

Bdnf and multiple CAMK components that transduce its signal are also H3K27me3-modified. BDNF signaling is a well-established regulator of GN radial migration. The knockout of *Bdnf* (*88, 89*) or of its downstream pathway components *CaMKKII* or *CaMKIV* (*90*) all cause GN migration defects. The opposite phenotype occurs in response to over-activation of the Bdnf pathway, decreasing the time to differentiation. Transfection of GNPs with a constitutively active CREB, the terminal transcriptional activator of BDNF signaling, accelerated terminal differentiation of GNPs in culture (*91*).

From a differentiation therapy perspective, not all differentiation genes are equal. The greatest interest lies with genes that could be activated to prevent subsequent re-entry into the cell cycle. Attempted cell division in differentiated neurons typically leads to cell death as seen with neurodegenerative disease (*92–95*) or to binucleated cells as seen with gangliogliomas (*96, 97*). Our present results suggest several candidate differentiation genes, but another approach is to manipulate Ezh2 directly and thereby control many of these genes simultaneously.

### Ezh2 inhibitors as a potential differentiation therapy

EZH2 inhibitors have been tested in SHH-subtype MB cells and have caused decreased cell viability (*98, 99*) and increased rates of differentiation (*100*). One rationale for using Ezh2 inhibitors in MB was that forced expression of *NeuroD1* in MB cells is capable of driving differentiation (*100*). ChIP qPCR analysis had demonstrated that *NeuroD1* is marked by H3K27me3, and Ezh2 inhibitors showed increased rates of differentiation both in culture and in vivo models. Therefore, EZH2 inhibition might derepress *NeuroD1* and spur differentiation of MB cells.

Our data provide genome-wide context for the previous work by identifying the large number of H3K27me3-modified genes during development and in MB cells. EZH2 drives H3K27me3-mediated repression in *NeuroD1* and in more than half of the 787 genes whose transcription increases in early GN differentiation. The mechanism by which EZH2 inhibition promotes cell death and differentiation of MB cells is likely to depend on its regulation of many genes in addition to repressing *NeuroD1*.

### Combined Ezh2 inhibition and cell cycle arrest as a differentiation therapy

A prevailing hope is that properly stimulated differentiation can override tumor cell proliferation, for MB and other cancer types (*101–103*). Differentiation therapy in acute promyelocytic leukemia using retinoic acid (RA), has dramatically extended survival of PML patients (*104, 105*). RA induces the cancer cells to differentiate from a granulocyte precursor into a mature myelocyte that can no longer divide (*106, 107*). Given its transformative effect in AML, RA was trialed for patients diagnosed with MB in hopes of similarly driving differentiation and long-term regression. Unfortunately, RA did not prove effective in MB and phase 2 trials were ultimately halted. RA does cause growth arrest of MB tumous (*108, 109*), but through apoptosis rather than induced differentiation (*110, 111*).

Our work suggests a nuanced relationship between proliferation and differentiation in GNPs and MB cells. Ezh2 appears to have a significant role in delaying activation of differentiation genes but *Ezh2* loss or inhibition is not capable overriding the cell cycle. *Ezh2* cKO did not reduce GNP proliferation (Fig 3) and there was only a marginal increase in differentiation with Ezh2 inhibition alone (Fig 8). Neuronal differentiation is a complex process and understanding where and how a given regulator fits into that process is critical to maximizing its use as a clinical differentiation therapy.

Additional observations raise concerns about using EZH2 inhibitors as a solo differentiation therapy. First, *Ezh2* cKO did not prevent MB formation in mouse models of MB. SmoM2 mice, who develop MB from unrestrained Hh signalling were crossed with Math1-Cre / Ezh2fl/fl developed MB at 100% penetrance as in normal SmoM2 control mice. The MBs from the *Ezh2* cKO mice were more aggressive than control SmoM2 with intact *Ezh2*, leading to early death of the mice (*78*). *Ezh2* cKO, in Ptch1^+/-^ mice also showed no impact on the rate of MB formation. The reduction in H3K27me3 with *Ezh2* cKO is substantial, exceeding what is possible with an Ezh2 inhibitor, so using Ezh2 inhibitors alone is unlikely to yield differentiation rates high enough to reduce cancer progression. Another concern is that *Ezh2* cKO has no apparent effect on proliferating GNPs. If anything, there were increased numbers of GNPs in G1 in *Ezh2* cKO mice compared to WT mice, indicating that GNPs will not cease dividing when H3K27me3 is strongly depleted. Because MB cells are dividing too, Ezh2 inhibition alone will probably not drive a high rate of differentiation.

With regards to forced G0 arrest in MB cells, our observations parallel long term animal experiments, where CDK4/6 inhibitors were used with xenografts of human MB cells. The tumors dramatically regressed when mice were treated with inhibitor (*112*), but three-quarters of the tumors recurred within 60 days of drug removal. This suggests that forced exit of the cell cycle is not sufficient to drive terminal differentiation of MB cells. In our experiments, wash-out of Ezh2 and CDK4/6 inhibitors resulted in cell-cycle re-entry of MB cells within 24 hours (Fig 8). When developmental regulation was closely recapitulated using dual inhibitors, driving both exit from the cell cycle and inhibition of Ezh2, we observed significantly higher levels of terminal differentiation without re-entry into the cell cycle following drug wash out.

### Barriers to clinical differentiation therapy

Our results suggest two critically important ideas for designing a differentiation therapy for MB. First, our evidence supports the use of Ezh2 inhibition as a strategy for the differentiation of Shh-subgroup medulloblastoma (*100*). The second is that premature transcriptional de-repression by Ezh2 inhibition is not sufficient to cause differentiation in a dividing MB cell. Prior to larger scale testing in animals, more extensive work needs to be done to optimize the duration of cell cycle arrest. A phase I trial of Palbociclib for pediatric brain tumours was recently completed which showed bone marrow suppression at higher doses (*113*). One advantage of differentiation therapy is that the cells may not need to be G0 arrested for more than 72 hours to get induction of differentiation, which means the CDK4/6 inhibitor could be delivered in short pulses. As bone marrow suppression took weeks to months to develop these short pulses will prevent accumulated toxicity (*114*). Continuous administration of an Ezh2 inhibitor would be supplemented with 72 hour cyclic period of CDK4/6 inhibition.

## Materials and Methods

### Crosslinking Chromatin immunoprecipiation (ChIP)

The ChIP protocol was adapted from an existing protocol (*115*). Briefly, a single cell suspension of GNPs or MB cells were crosslinked with 1% formaldehyde for 10 min, then quenched with 2.5 M Glycine. Chromatin was sheared using a Bioruptor (Diagenode, Denville, New Jersey) for 6 cycles of 15 minutes (30 sec on, 30 seconds off at maximum power).

Following sonication the insoluble material was pelleted. Antibodies were added to the sheared chromatin as indicated by table 1 to 750 ul of chromatin and incubated at 4 C overnight. For each ChIP-seq experiment 3 technical replicates of the immunoprecipitation were performed and pooled after DNA isolation. Protein G agarose beads (Roche) were used to precipitate the antibody bound protein and 4 sequential washes were done with the buffers described in the original paper (*115*). Protein was eluted of beads with 10 mM EDTA and 1% SDS and the DNA was liberated from protein using Proteinase K (Roche). The DNA was purified using phenol chloroform extraction then treated with RNAse A. DNA was quantified using Qubit DNA HS (Invitrogen). ChIP-seq libraries were prepared using NEBNext ChIP-Seq Library Prep (New England Biolabs). Sequencing was performed using HiSeq 2500 (Illumina Inc) with 40 BP single end reads.

**Table 1:** Antibodies and concentrations used for ChIP.

| Antibody | Vendor | Species | Serotype | Catalog | Amount<br>(ug, ul)<br>in 750 ul<br>lysate | RRID |
| --- | --- | --- | --- | --- | --- | --- |
| H3K4me3 | Abcam | Rabbit<br>polyclonal | IgG | ab8580 | 5 ug | AB_306649 |
| H3K4me1 | Abcam | Rabbit<br>polyclonal | IgG | ab8895 | 5 ug | AB_306847 |
| H3K27me3 | Milipore | Rabbit<br>polyclonal | IgG | 07-449 | 5 ug | AB_310624 |
| H3K27ac | Abcam | Rabbit<br>polyclonal | IgG | ab4729 | 5 ug | AB_2118291 |
| Ring1b | Active<br>Motif | Rabbit<br>monoclonal | IgG | 39664 | 10 ul | AB_2615006 |
| H2Aubi119 | Milipore<br>(Upstate) | Mouse<br>monoclonal | IgM | 05-678 | 10 ug | AB_309899 |
| H3K36me3 | Abcam | Rabbit<br>polyclonal | IgG | ab9050 | 5 ug | AB_306966 |
| RNA<br>polymerase<br>2 (S5<br>phospho) | Abcam | Rabbit<br>polyclonal | IgG | ab5131 | 5 ug | AB_449369 |
| RNA<br>polymerase<br>2 (S5<br>phospho) | Active<br>Motif | Mouse<br>monoclonal | IgG | 39097 | 10 ul | AB_2732926 |

**Table 2:** Mice alleles used.

| Allele | RRID |
| --- | --- |
| Math1-Cre | IMSR_JAX:011104 |
| *Ezh2*<sup>flox</sup> | MMRRC_015499-UNC |
| Math1-GFP | MGI:4456122 |
| *Ptch1*<sup>+/-</sup> | IMSR_JAX:003081 |

### Native ChIP

The crosslinking protocol described above was performed for all antibodies in Table 1, except the H2Aubi119 antibody where native ChIP was used. The protocol for native ChIP has previously been described (Hasegawa et al., 2016). In brief, cell lysate from fresh frozen GNPs or MB was incubated in MNase for 20 min at 4°C and stopped by adding 1/25 0.5 M EDTA. The amount of MNase used for each reaction was batch adjusted to obtain mono-nucleosomes after a 20 min digestion. After centrifugation at 13000 rpm for 15 min at 4C, the supernatant containing the mononucleosome fraction was collected. Protein was precipitated using Protein G Dynabeads (Invitrogen), with rabbit anti-IgM IgG linker antibodies for the H2Aubi119 ChIP. The beads were washed 3 times using buffers described in Hasegawa et al (2016). DNA was eluted, freed from protein using Proteinase K (Roche), and purified using phenol chloroform extraction. Preparation of ChIP-seq libraries was performed as described for crosslinking ChIP.

### ChIP-seq data processing

Reads were adaptor filtered, trimmed, mapped to mm9 using Bowtie2. From the mapped reads duplicates and reads with a mapping quality below 20 were discarded. Quantification of a region around the TSS which were defined in the previous section proceeded with the following steps. Peaks in aligned reads for histone modifications were called using EPIC (https://github.com/biocore-ntnu/epic), which is a more computationally efficient version of SICER (*116*). Macs2 (*117*) was used to call peaks for the ATAC-seq data as no input sequencing was done. Many of the histone modifications would vary substantially in terms of both the number of reads in a set genomic region but also in terms of the width of the marked area. To capture the variation in with the promoter modifications were quantified, for each of the annotated transcriptional start sites (TSSs) with a window of 250 base pairs flanking the site was quantified and extended to include overlapping peaks. All quantified promoter features underwent a regularized log transform (*118*) and were scaled 0 to 1.

For RNAseq and H3K36me3 we altered our quantification strategy. H3K36me3 is a histone modification associated with an elongating RNA polymerase II seem during active expression, which is absent over the promoter. The H3K36me3 ChIP-seq was quantified across the entire gene body and then divided by the length of the gene. The P7 RNAseq data was quantified across all exons and divided by the total length of the exons. Both RNAseq and H3K36me3 were then scaled between 0 and 1.

To calculate the enrichment of chromatin modifications, epigenetic features were separated based upon the distribution of the quantified modifications. The normalized data show a clearly bimodal distribution or a distribution with a large upper tail. Marked or bound versus unmarked or unbound genes could be distinguished by fitting two normal distributions. First long upper tails were removed by calculating the first order derivative of the smoothed ranked data similar to how super enhancers are identified (*42*). We set the threshold for the long upper tails at 0.0004 pragmatically. The remainder of the datapoints for each tracks were split using a mixture of Gaussian distributions using mixtools (*119*). Each data set were split into marked/unmarked/tail for the enrichment calculation and for plotting.

### RNA-seq

RNA was extracted using Trizol (Invitrogen), with and mechanical disruption with a 1.5 ul tube mortise and pestle. Following precipitation, the RNA was treated with DNase1 and cleaned up with a RNAeasy colomn (Qiagen). RNA was quantified with the Qubit and the quality of the RNA was assayed using Agilent Bioanalyzer. Libraries were prepared using mRNA purified using polyA bead selection (New England Biolabs), using Illumina TruSeq v2 library preparation kit (Illumina). Sequencing was performed using HiSeq 2500 with 100 BP single-end to approximately 60-70 million reads per sample.

### Generating cell type-specific transcript models

Since we were attempting to identify epigenetic regulatory features by looking for common signatures present in a large number of genes, it was important that each protein coding gene would be counted roughly equally. Some genes have numerous annotated TSS while others only have one and the discrepancy could bias the downstream analysis. We wanted to represent each protein coding gene with at least one TSS and one transcript model but permitted a gene to have multiple TSS and associated transcript models if there was evidence in the ChIP-seq data of an additional TSS. We used both the Ensembl mouse 67 annotation system as well as the UCSC mm9 knownGene annotation to generate a robust list of baseline TSSs and transcript models. The TSS in the reference annotation was quantified using both the GNP H3K4me3 data and the adult cerebellum H3K4me3 data from the Encode project (*120*).

For genes with clear H3K4me3 binding we selected transcripts with the strongest H3K4me3 at the TSS or within 500 BP of the optimal TSS. For genes without clear H3K4me3 over any transcript we took the maximum of all of the other modifications annotated over potential TSS. To identify the best transcript model for each gene from the numerous in the reference transcriptome we performed de-novo transcript assembly from 1 replicate in each of the time points in the RNA-seq data using Stringtie (*121*). The time point transcript assemblies were then merged using Stringtie merge without the reference annotation. Both selecting a TSS and transcript model using only de novo assembly is complicated by a number of issues. The PolyA selection is 3’ biased, which will bias the de novo transcript models towards TSS which occur in the 3’ direction in a manner that is worse for low expressed genes. We calculated an overlap for each transcript between the reference transcript library (UCSC and ensemble) and the Stringtie models using Bedtools (*122*). We then selected a transcript model from the reference with the highest overlap to the Stringtie models for each gene.

### RNAseq quantification

The RNAseq data was mapped to mm9 using Tophat2 (*123*), so the last mm9 associated Ensembl release (release 67) was used as reference transcriptome. The transcript models used to quantify the RNAseq data developed as described above. Transcripts were quantified with HTSEQ. The overall pipeline was developed with modification from the Bradner lab pipeline (https://github.com/BradnerLab/pipeline).

### Processing existing data sets

The Hatten lab NeuroD1-TRAP data was downloaded from GEO (GSE74400) and the Affemetrix annotations linked to the Ensembl mouse 67 transcript models were downloaded using biomaRt. Initial analysis was performed using oligo and limma packages from Bioconductor. The probes were then collapsed to the transcript models described above.

The scRNAseq data from the Taylor lab was obtained from GEO (*31*) and the reads were processed using the DropletUtils package from Bioconductor. We used the annotated cluster to cell tables provided in the supplementary data to reconstruct the data. The granule neuron lineages were separated from other cell types like Purkinje cells, astrocytes. In order to eliminate genes from the bulk RNAseq that were likely arising from non-granule neuron lineage we used the existing cell type clustering analysis done by the Taylor lab using Seurat. The cell types were separated into granule neuron lineage and non GN lineage. For each gene, across the cell types, the highest expression in GN lineage and the non GN lineage cells was calculated. By comparing the maximum GN expression to non GN expression we excluded genes with the following 3 criteria. We excluded genes that were expressed in alternative lineages when not expressed at all in the GN lineage cells, genes that were 2 fold higher in non GN or when they were significantly higher expressed in non GN lineage by the Taylor lab Seurat analysis.

### Differentiation gene classification

Expression cutoffs for transcriptional time course data were established using the distribution of the maximal values from the time course. For each dataset the average maximum of the 3 highest expressed timepoints/replicates were taken. Based upon the distribution we created a mixture model to estimate the point at which the probability of the expressed versus unexpressed distribution was equal to .50. For the Scott lab RNAseq dataset the count data was highly zero weighted, so we utilized a mixture of 2 gamma distributions (*124*). The Hatten lab TRAP data was modeled using 2 normal distributions. The transcriptional profiles were merged by interpolating the overlapping time points. We then calculated the timepoint that had the maximum and minimum RNA using the combined dataset. Using the maximum and minimum values we calculated a fold change using both datasets. We also estimated the developing time point at which the gene reached 50% of its maximum transcript abundance (t50). In order to be accepted as a significant changer p-values were calculated based upon each dataset individually, using DEseq2 (*118*) for the RNAseq dataset and limma (*125*) for the TRAP data.

### Mouse husbandry

Ezh2 conditional knockout (Math1^Cre;^ *Ezh2^flox^*^/flox^) mice were generated by crossing Math1>Cre (*56*) with the *Ezh2^flox^*^/flox^ mouse where Loxp sites flank the catalytic SET domain of Ezh2 (*55*). Mice were then bred homozygous for the flox Ezh2 allele, and Cre negative littermates were used for controls. Math1>GFP (*30*) mice were maintained as homozygous.

Math1^Cre;^ *Ezh2^flox^*^/flox^;Math1>GFP mice All animal studies were approved by the Stanford APLAC review board protocol.

### Spontaneous MB tumor formation

ChIPseq and RNAseq data was generated from spontaneous MBs that occurred in Ptch1^+/-^ mice, with MB formation determined by following for neurologic decline. Since the generation of MBs can have severe neurological affects, animals were monitored daily for physical abnormalities such as ataxia, hunching, immobility. Math1^Cre;^ *Ezh2^flox^*^/flox^;Ptch1^+/-^ and *Ezh2^flox^*^/flox^;Ptch1^+/-^ mice were also generated to test the effect of *Ezh2* cKO on MB formation.

### Edu pulse/chase

To assess migration of the GNPs out of the EGL we performed 48 h Edu pulse chase. A single IP injection (50mg/kg using a 5mg/mL stock diluted in PBS) was administered to p5 *Ezh2* cKO and wild-type mice. After 48 h, brains were dissected and fixed in 4% PFA overnight then transferred into 30% sucrose for 24h. Fixed whole cerebellum were mounted in OCT and sectioned at 20μm. EdU staining was performed as per manufacturers instructions (Life Technologies, Clik-iT Plus EdU Alexa Fluor 647 Imaging Kit, cat no. C10640). Following Edu labeling, sections were blocked in 0.2% triton X-100 and 5% Donkey serum 1h at RT. Sections were then stained with Rabbit anti-p27 as described in the tissue immunofluorescence section.

### GNP isolation and purification

The GNP extraction and purification has been previously described (*1, 126*). In brief, the mice were euthanized and the cerebella were dissected and minced. The cerebellar pieces were incubated with 10 U/ml papain (Worthington, NJ, United States, LSOO3126) and 250 U/ml DNase (Sigma, MO, United States, D4627) in HBSS (Stem Cell Technologies Canada Inc, Vancouver, Canada, 37150) at 37 C for 30 min and the papain was halted using 8 mg/ml Ovomucoid (Sigma, MO, United States, T2011) and 8 mg/ml bovine serum albumin (Sigma).

The pieces were then sequentially triturated in the presence of DNase to prevent clumping, then passed through a 70uM nylon cell strainer (VWR, 21,008-952). GNPs were isolated using a 35%, 65% Percoll (Sigma, P4937) step gradient with centrifugation at 2500 rpm for 15 min.

### E15.5 GNP purification

Because of the small cell number and the contamination of Sox2 + cells we utilized FACS to purify the pre-Shh GNPs. Timed pregnant mothers homozygous for the *Math1*-GFP reporter were sacrificed using cervical dislocation following deep anesthesia. The embryos were harvested, and the rhombic lip and cerebellar anlage were dissected using micro-scissors and jeweller forceps. The single cell suspension was generated as above. GFP+ were isolated using standard FACS technique. The cells were then lyzed in Trizol without centrifugation and RNA purification proceeded immediately.

### MB cell culture

MB cell lines (*61*) generated from *Ptch1*^+/-^ mice by the Seghal Lab (Dana-Farber Cancer Institute, Harvard Medical School), and *Ptch1*^+/−^; *Tpr53*^−/−^ from the Rudin Lab (Memorial Sloan Kettering Cancer Center), were grown as suspension as neurospheres. Briefly, cells were maintained in DMEM/F12 supplemented with glutamine and with B27 minus vitamin A, penicillin and streptomycin. The cells were passaged every 6 days by collecting the spheres by gravity, washing, followed by dissociation with Accutase (Stemcell Technologies 07920) and replating as a suspension to allow cells to spontaneously re-aggregate. UNC1999 was soluble in DMSO while Palbociclib was soluble in water (Selleckchem).

### GNP culture

P7 GNPs were purified as described above. The cerebella dissected either from the *Ezh2* cKO described above or from mice with a Math1>GFP (*30*) reporter that delineates transit amplifying GNPs from differentiated cells. The GNPs were grown as described previously for our GNP proliferation conditions (*1*) with 3ng/ml of Shh 461-54 (R&D systems). The cells were fixed at 24 h and immunofluorescence imaging occurred. To measure the protein abundance of the product from the differentiation genes, the freshly isolated cells were plated without Shh to induce differentiation, fixed after 6h, 24h or 48h in culture.

### Cell culture imaging

All cell culture imaging for both GNPs and MB cells was performed on 96 well, glass bottom plates (Cellvis P96-1.5H-N). The plates were coated with 100 ug/ml PDL (Millipore A-003-E) for 3 h at 37 C, then washed, and coated with 10 ug/ml Lamnin (Millipore CC095) overnight at 37 C.

GNPs were purified as described above and MB cells were dissociated with Accutase as described above, then filtered using a 70-micron to remove clusters of cells. Following cell counting 10,000 cells were plated per well and perturbations were performed 2h after plating. Following treatment, the cells were fixed with 4% paraformaldehyde for 10 min at room temperature. The cells were then blocked using 5% donkey serum, 1% BSA and 0.2% triton X-100 for 1h at room temperature. Primary antibodies were incubated overnight at 4 C. The primary antibodies used are described in table 3. Cells were co-stained with Rhodamine Phalloidin (Molecular Probes R415). Donkey anti IgG secondary antibodies against mouse and rabbit were used conjugated to Alexa-488 and Alexa-647 at 1:500 (Jackson Immunoresearch). All cell imaging was performed using the ImageXpress Micro XLS Widefield High Content Screening System (Molecular Devices, Sunnyvale, CA) using 20x (0.75 NA) Nikon objectives. The intensity of fluorescence in each cell was automatically calculated using custom MATLAB scripts. Nuclei were segmented using DAPI as previously described (*127*). Downstream analysis was performed in R.

**Table 3:** Antibodies used for immunofluorescence.

| Antibody | Vendor | Species | Serotype | Catalog | Dilution | RRID |
| --- | --- | --- | --- | --- | --- | --- |
| NeuN | Millipore | mouse monoclonal | IgG | MAB377 | 1:200 | AB_2298772 |
| Phospho-Rb (Ser807/811) clone D20B12 | Cell Signaling Technology | rabbit monoclonal | IgG | 8516 | 1:1000 | AB_11178658 |
| p27 | Abcam | Rabbit monoclonal | IgG | ab32034 | 1:100 | AB_2244732 |
| Cbx7 | Abcam | Rabbit polyclonal | IgG | ab21873 | 1:200 | AB_726005 |
| H3K27me3 | Active Motif | Rabbit polyclonal | IgG | 39155 | 1:200 | AB_2561020 |
| Map2 | Abcam | Chicken polyclonal | IgG | ab5392 | 1:2500 | AB_2138153 |
| Ezh2 clone D2C9 | Cell Signaling Technology | Rabbit monoclonal | IgG | 5246 | 1:200 | AB_10694683 |

### Tissue Immunofluorescence

P7 mice were perfused with 4% PFA, then fixed for 16 h. Fixed whole cerebellum were mounted in OCT and sectioned at 20μm. Antigen retrieval was performed only for Cbx7 using 10 mM Citrate pH 6 buffer, boiled for 15 min. Immunofluorescence proceeded as described in the cell culture staining protocol. Panoramic Imaging was done on a Zeiss Axioimager and 40x confocal micrographs were generated using a Leica Sp2 confocal microscope.

### Cerebellar layer segmentation

To segment the cerebellum into outer EGL, inner EGL, ML, IGL and deep white matter we generated panoramic tiled immunofluorescence images of the P7 cerebellum, with DAPI, NeuN (Alexa 488) and p27 (Alexa 647). The images were then background corrected and stitched in Fiji (Image J) using Grid/Collection with 20% overlap. A custom MATLAB script was then used to perform the segmentation. The first pass performed preprocessing on the image and coarsely identified each layer using a combination or ratio of the individual channels. The IGL was defined as NeuN high, the inner EGL as p27 high, and the outer EGL as DAPI high. Each layer of the cerebellum borders the following layer, allowing previously established layers to serve as guides for the development of following layers. Beginning from the inner most layer, through to the outermost, dilations were used to define and fill out the regions of interest. The pixels of each region were filtered and smoothed to promote clear boundaries between the layers. A composite image was then produced containing all the regions. With each region clearly defined, calculations for segmented pixel area and region measurements for NeuN, P27, and DAPI fluorescence intensity for each layer could be determined.

### Cerebellar immunofluorescence quantification

The images were generated as described above. Linear segments centered along the pia, with an EGL on either side were selected and made into separate images. The image segments were further subdivided into 200 pixel sections moving perpendicular to the axis of the EGL. A line track was generated by taking the pixel mean perpendicular to the EGL. The individual line segments were then aligned using cross-correlation from the Stats package in R. After alignment the symmetric line segment was split by calculating the local minimum near the center of the dataset for an average p27 intensity of the entire dataset. The center was set to 0 on the x-axis which allowed us to split each line segment into 2 moving from 0 the pial boundary to the IGL at the highest values. There was variability in the distance from the center point to the pial boundary where the EGL starts caused by a gap in the arachnoid along the invagination of the folia. The pial boundary was determined as a drop off in the immunofluorescence background. To find this boundary we identified the peak change in the product of NeuN and p27 values.

### Quantification of cell cycle and differentiation and process extension

Dead cells and G2 cells were separated from G0/1 cells using the cell pixel area and DAPI median values, then G0, G1 and G2 were further separated according to the level of Ki67 or RB phosphorylation (Ser807/811). Differentiated cells were identified using a cut-off value for NeuN immunofluorescence only accepting differentiating cells if they are marked as G0 by the above-described criteria. Process extension and process length was quantified using Neuroncyto 2 (*60*) image by image using Map2 immunofluorescence with DAPI labeling of the nuclei.

## Additional information

### Data availability and resource sharing

Data supporting this study are available in GEO GSE279346 and GSE279347. Custom analysis scripts are available following GITHUB repositories; jpurzner/seq_pipelines, jpurzner/chip_tools, jpurzner/cell_culture_segment, jpurzner/layer_quant, jpurzner/cerebellar_segmentation.

## Acknowledgements

The research presented here was funded by the National Institute Health (5R01CA157895-02), the American Brain Tumour Association, B*CURED and the American Association of Neurologic Surgeons via the Neurosurgery Research and Education Fund.

## Author contributions

JP, ASB, TP, KW, MDT, YJT, MTF, MPS designed research. JP, ASB, TP, LE, SB, UL, AK, KW performed research. JP, ASB, TP, KA, AS, KW, MDT, YJT, MTF, MPS analyzed data. JP, TP, ASB and MPS wrote the paper.

