## Supplemental Figures for "Ezh2 Delays Activation of Differentiation Genes During Normal Cerebellar Granule Neuron Development and in Medulloblastoma"

**Supplemental Fig 1**

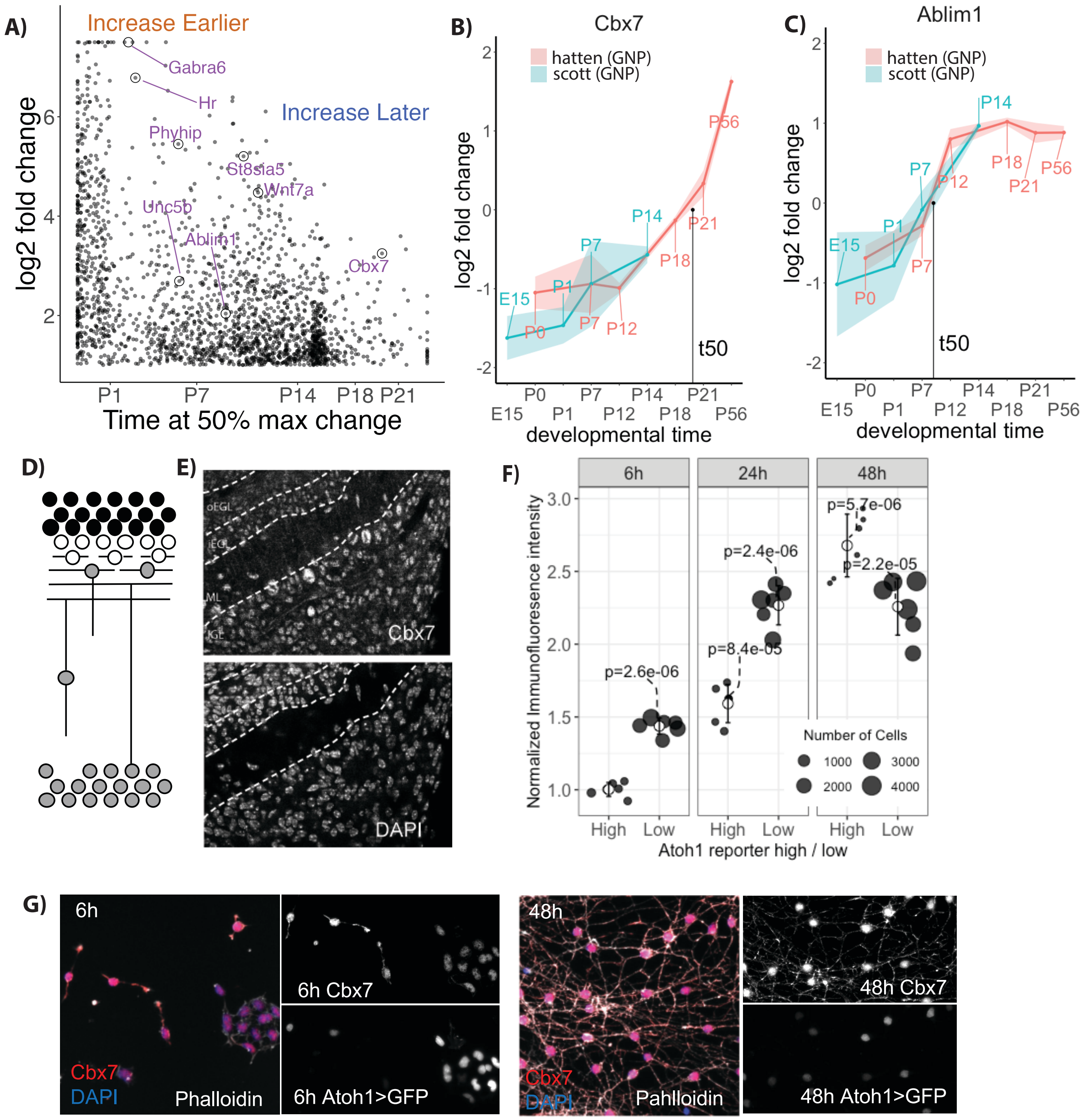

**Supplemental Figure 1: Validating measured transcription changes by looking at changes in protein abundance during GNP differentiation.**

**A)** Plot of the time when a differentiation gene reached 50% of its maximal expression (t50), by the combined fold change of both datasets. Example differentiation genes are circled, demonstrating diverse timing for transcriptional changes during granule neuron differentiation. **B)** Interpolated transcriptional time course of RNA abundance during developmental time from E15 to P56 (turquoise line = our RNASeq, red line = Hatten NeuroD1-TRAP) for the differentiation genes *Cbx7,* which increases later and **C)** *Ablim1* which increases earlier. **D)** Schematic of the P7 sagittal cerebellar slice showing from outside in; outer EGL (dividing), inner EGL (cell cycle exit), ML (migration) and IGL. **E)** Validation of transcriptional data with Cbx7 protein abundance in sagittal sections of P7 cerebellum, showing increased Cbx7 in the IGL (DAPI=nuclear marker). **F)** Quantification of cultured GNPs from Math1>GFP mice, labelled with Anti-Cbx7 over time show an increase in Cbx7 as GNPs differentiate. Relative fluorescent intensity of Cbx7 in Atoh1 high (proliferating GNPs to early differentiation) and Atoh1 low (differentiated) GNPs (n = 6). P-values from paired t-test with each group against the 6h Atoh1 high group**.** **G)** Examples images of the cultured GNPs that were quantified in F.

#### Supplemental Fig 2

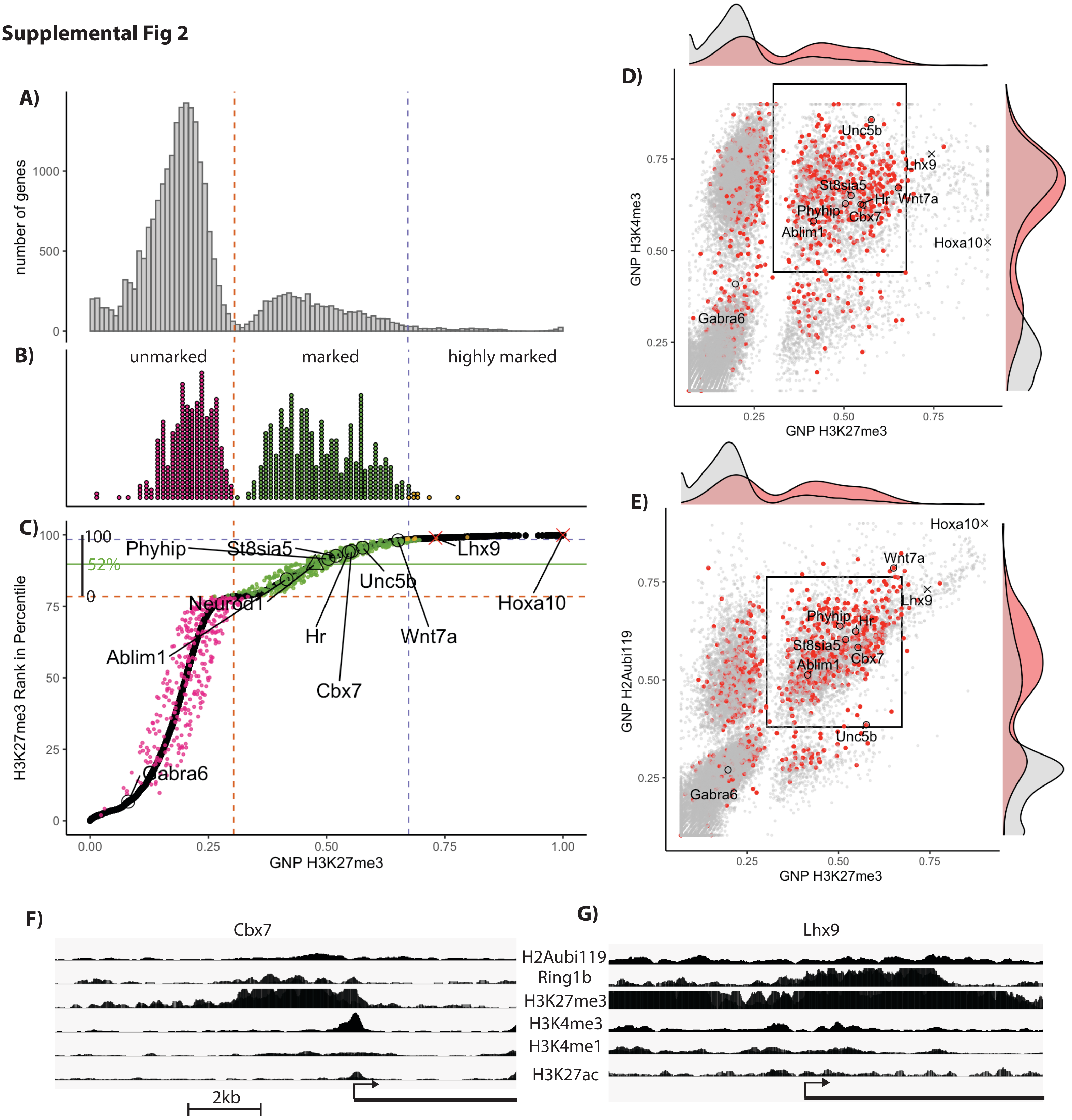

**Supplemental Figure 2**: **H3K27me3 modified differentiation genes show moderate amounts of H3K27me3, H2Aubi119 and are also marked with H3K4me3.**

**A)** A histogram showing the H3K27me3 ChIPseq from P7 GNPs, quantified over all genes. The x-axis shows the normalized H3K27me3 for each promoter region as described in the Materials and Methods. The orange dotted line separates genes that are unmarked vs marked by H3K27me3 in in GNPs. The blue dotted line separates main body of bound genes from a long tail of genes associated with very high levels of H3K27me3. **B)** A histogram of putative differentiation genes showing more genes within the bound region (green) compared to the unbound (purple) or the long tail (yellow). **C)** A rank plot showing the cumulative percentile of all genes ranked in order of H3K27me3 level (y-axis) by normalized H3K27me3 (x-axis) with the differentiation genes labeled as above. The y-axis includes a secondary bar that shows the percentile rank of just the bound group. The green line at 52% shows the mean value of the bound percentile rank of the differentiation genes indicating a moderate amount of H3K27me3. **D)** Scatter plot of H3K27me3 and H3K4me3 with the differentiation genes marked in red. A box is drawn around the center of the H3K27me3 marked differentiation genes. Marginal histograms are shown with the red density plot being the differentiation genes. Examples differentiation genes are marked with a black circle and comparative genes that are more heavily repressed are marked with an X **E)** Similar scatter plot showing H2Aubi119 and H3K27me3, the differentiation genes are again red. **F)** A sequencing read pile up image showing the amount of ChIP-seq reads near the promoter for *Cbx7* and **G)** *Lhx9*. There is considerably more H3K27me3, Ring1b and H2Aubi119 at the heavily repressed *Lhx9*. The x-axis is base pairs across the gene promoter and the y-axis is number of reads by ChiP-seq.

### Supplemental Figure 3

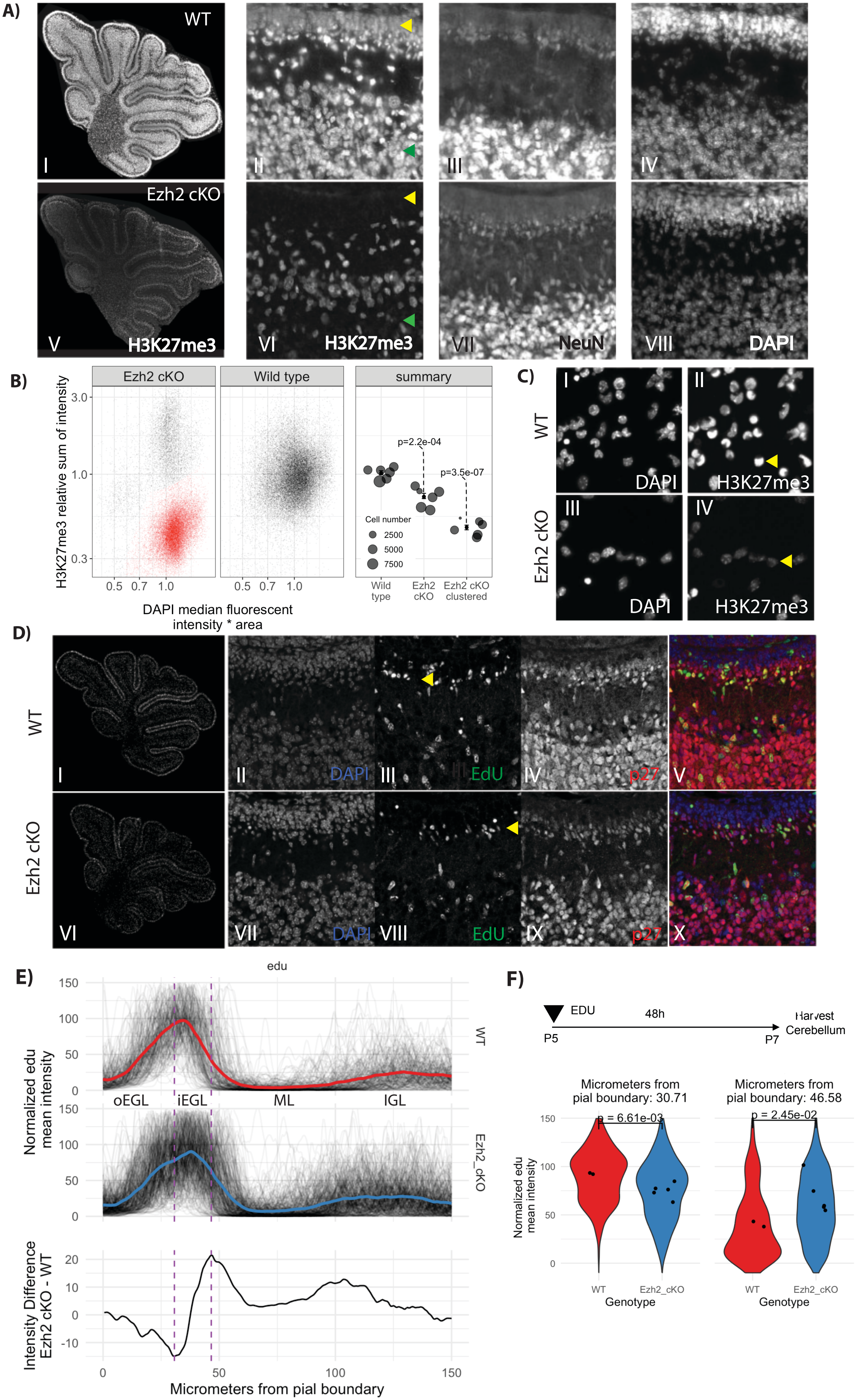

**Supplemental Figure 3: The sparse inner EGL is due to early nuclear migration, not to a defect in migration:**

**A)** H3K27me3 immunofluorescence of *Ezh2* cKO versus control in whole cerebellar sections, with enlargement of the same image showing HK27me3 (panel I, II, V, VI) , NeuN (III, VII), and DAPI (IV, VIII)**.** Comparison of panel II and VI show substantial drop in H3K27me3 in the EGL (yellow arrow) and IGL (green arrow) in the mutant. **B)** Single-cell quantification in cultured GNPs from *Ezh2* cKO and WT mice of H3K27me3 immunofluorescence showing a substantial drop in labeling in the majority of cells. Quantification includes the entire Ezh2 cKO group of cells or the GNP lineage cells only (labeled in red) which was separated using gaussian classification, both showing a statistical difference as measured by T-test (n = 6 per group). **C)** Example of cultured GNPs from Ezh2 cKO (panel III, IV) and WT (I, II) mice for H3K27me3 (II, IV) and DAPI (I, III), shows higher H3K27me3 immunofluorescence in WT (II) compared to Ezh2 cKO (IV). A representative GNP nucleus is shown with a yellow arrow in pane II and IV. **D)** To measure how far a given GNP had migrated since its last S phase we performed a EDU pulse-chase experiment with a 48 h delay. Whole cerebellum 20x panoramic reconstructions for WT (I) and *Ezh2* cKO (VI) show a less defined inner EGL across the entire cerebellum with *Ezh2* cKO. Confocal micrographs of DAPI (II, VII) , EDU (I, III, VI, VIII) and p27 (IV, IX) show in WT the 48h pulse labeled cells have come to rest along the border of the inner EGL, while in the *Ezh2* cKO labeled cells are seen entering the molecular layer (ML) as seen with the yellow arrow. **E)** Quantification of EDU from the pial boundary to the IGL in WT and *Ezh2* cKO along with the average difference between WT and *Ezh2* cKO. **F)** Violin plots showing that EDU labeling was more prominent further into the ML in the *Ezh2* cKO (n = 5) then the WT (n = 2) .  The P-value was determined by T-test.

### Supplemental Figure 4

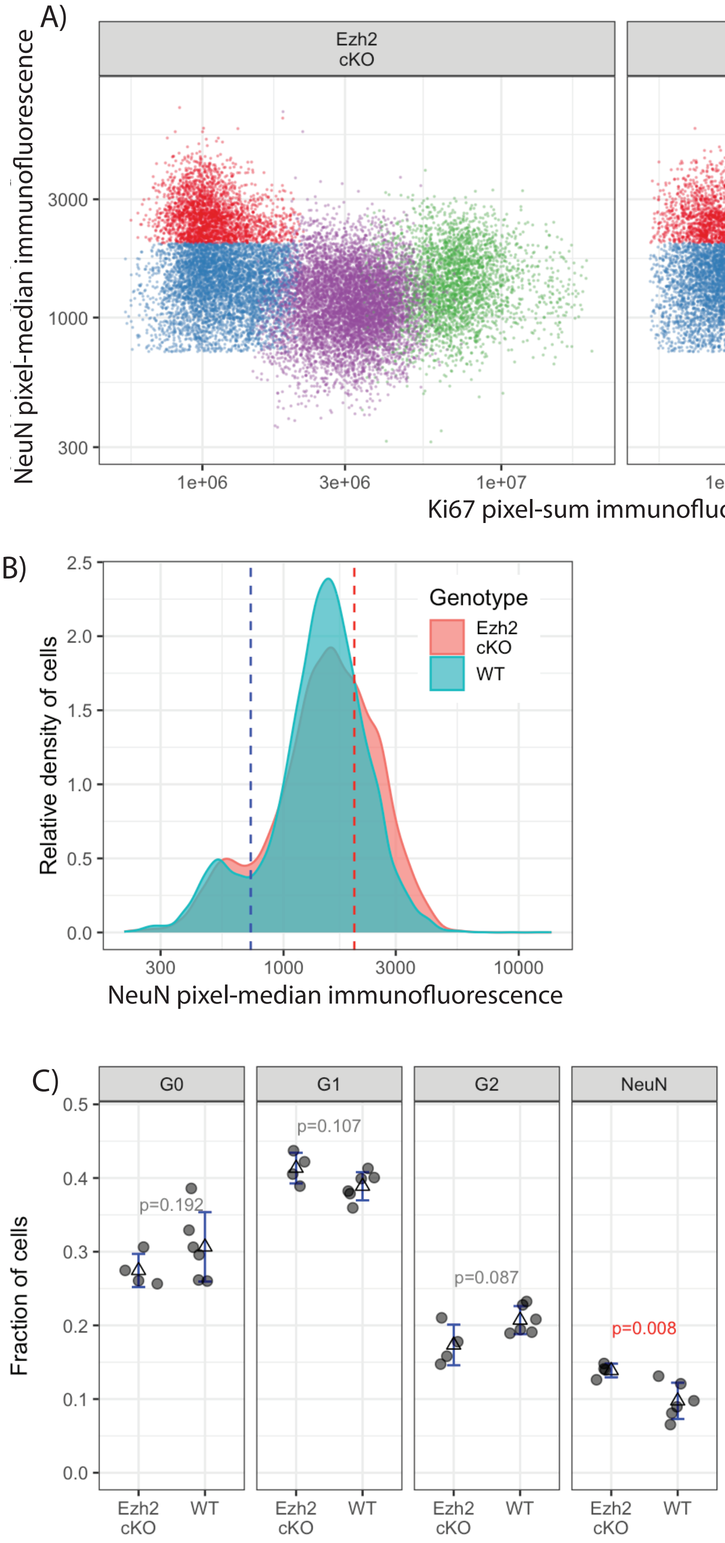

**Supplemental Figure 4: Ezh2 cKO leads increased differentiation in cultured cells by NeuN immunofluorescence**

GNPs cultured with Shh followed by labeling with Ki67, DAPI, and NeuN to measure cell cycle and differentiation in *Ezh2* cKO and WT. **A)** NeuN classification along with cell cycle characterization in a scatter plot of Ki67 and NeuN. G1/G2 distinction was established using DAPI. **B)** NeuN immunofluorescence histogram with labels for cutoffs for low and high expression. Both **A)** and **B)** shows an increased number of NeuN + cells in the *Ezh2* cKO. **C**) The fraction of cells within each class, demonstrating Ezh2 cKO (n = 4) had a higher fraction of NeuN+ differentiated cells then WT (n = 6). The p-value calculated by T-test. There was no significant difference in the number of cells In G1 or G2.

**Supplemental Figure 6**

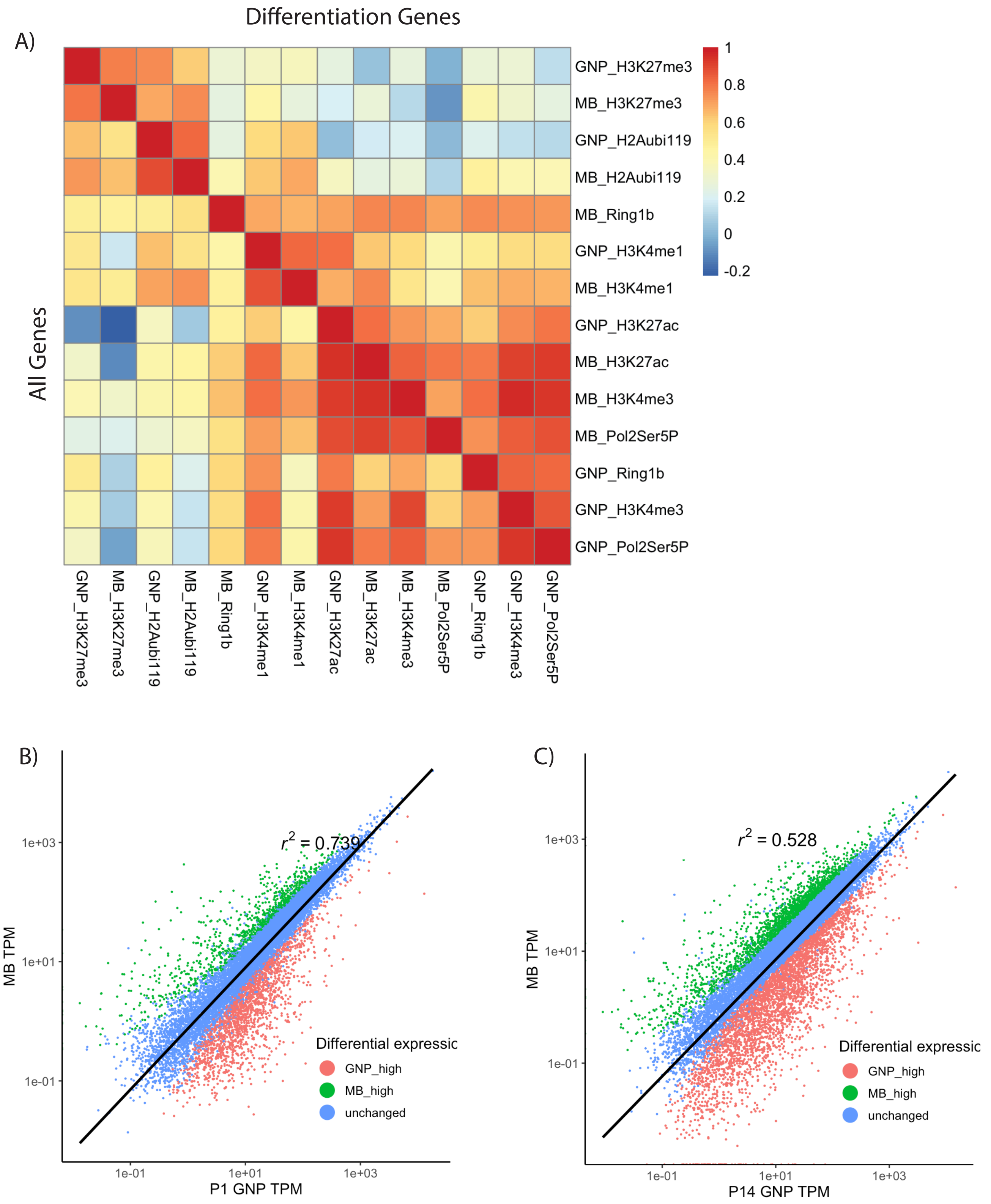

**Supplemental Figure 6**: **Chromatin modifications near promoter regions in GNPs s are highly correlated with those in *Ptch1*+/- MB cells.**

**A)** Pearson correlation values for GNP and MB chromatin marks, according to the color scale at right. The upper diagonal represents only differentiation genes while below the diagonal all genes are represented. **B)** Scatter plots of gene expression in MB compared to P1 and **C)** P14. Color code shows 2 fold higher expression in MB (green) or lower in MB expression (red) and adjusted p-value $<0.05$. The $R^2$ values are shown within panel B and C, were lower then for P7 GNPs (scatter plot in Fig 6B) which was $R^2 = 0.796$ indicating MB are most similar to P7 GNPs.

**Supplemental Figure 7**

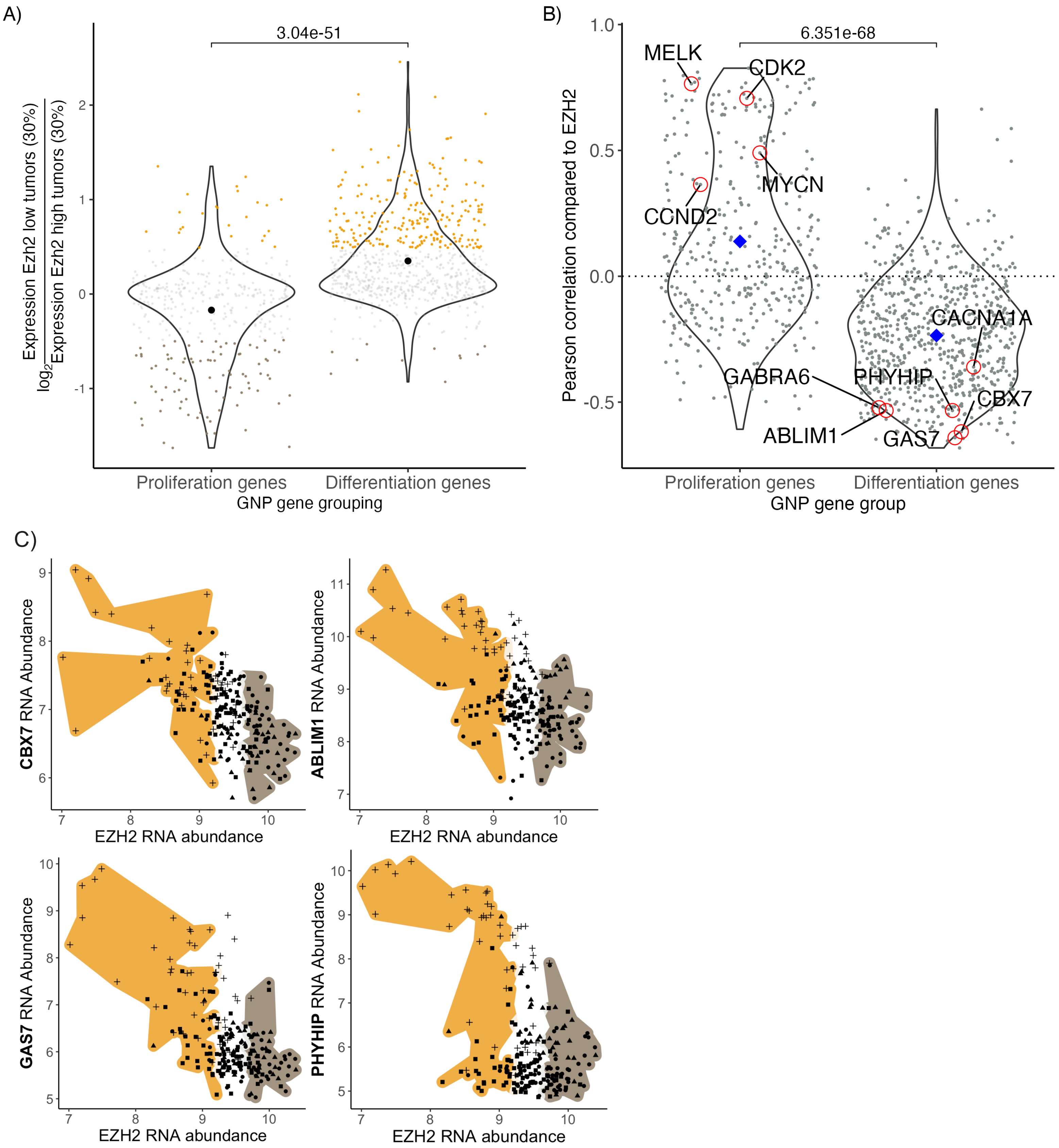

**Supplemental Figure 7**: **GNP differentiation genes anti-correlate with *EZH2* expression in human SHH-type medulloblastoma.**

**A)** On average the differentiation genes are 1.3 fold higher expressed in *EZH2* low MBs. A jitter plot showing the fold change in the differentiation and proliferation genes. The p-value was calculated with t-test. **B)** The expression of the differentiation genes are predominantly anticorrelated with EZH2 expression as seen with a jitter plot of the Pearson correlation co-efficient. The differentiation genes show a negative Pearson co-efficient with an average of -0.24 as indicated with the blue diamond. The p-value was calculated with the Wilcoxon rank-sum test. **C)** Plot of a selected differentiation CBX7, ABLIM1, GAS7, and PHYHIP versus EZH2 transcript abundance. Each dot is a human tumor. The shape indicating SHH subtype and color labelling are the same as in panel A. The Pearson correlation co-efficient for CBX7, ABLIM1, GAS7, and PHYHIP were respectively -0.61, -0.53, -0.64 and -0.53.

**Supplementary figure 8**

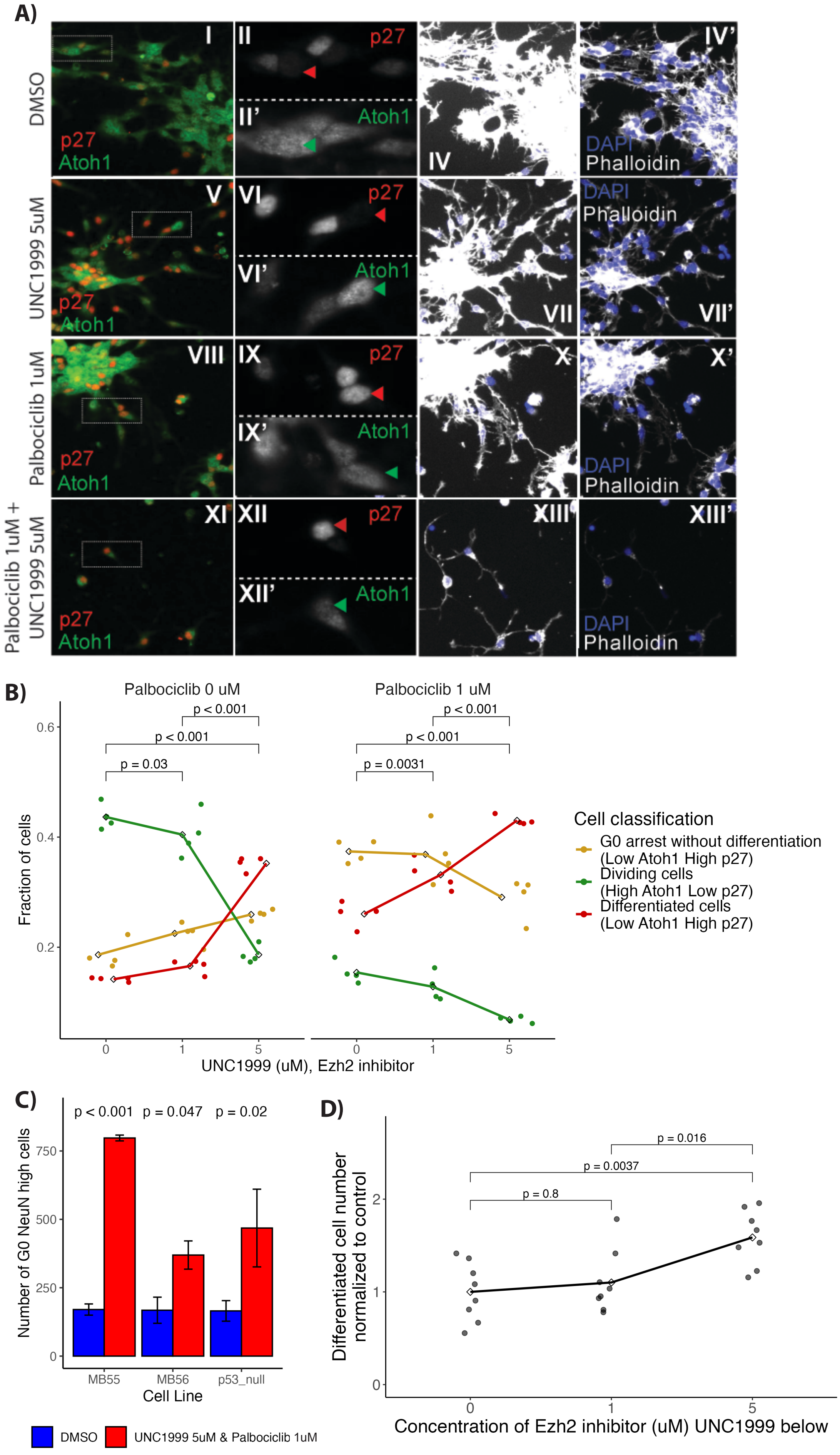

**Supplemental Figure 8: Combined treatment of MB cells with Ezh2 and CDK4/6 inhibitors increases markers of neuronal differentiation.**

A**)** Alternative quantification of differentiation in MB cells with labels for p27(red), Atoh1 (green), DAPI (blue) and phalloidin (white) levels in cells treated with DMSO (i-iv), Ezh2 inhibitor UNC1999 5 uM, (v-vii) CDK4/6 inhibitor Palbociclib 1 uM, (viii-x) dual treatment with Palbociclib 1 uM and UNC1999 5 uM. (xi-xiii) without washout. The dashed boxes in i, v, viii and xi show an enlargement of cells in ii and vi respectively, the individual colour channels are seen as separate panes. The red arrows indicate p27 + cells and the green arrows indicate Atoh1 cells. **B)** Quantification of fractions of cells in the following groups: Differentiated cells (red) that have high p27 and low Atoh1, G1/2 cells have low p27 and high Atoh1 (green), transient G0 cells have high p27 and high Atoh1 (golden). The highest rates of differentiation occurred with combined Palbociclib 1 uM and UNC1999 5 uM. P-values comparing the fraction of differentiate cells were calculated using ANOVA with post-hoc Tukey test (n =4). **C**) The absolute number of differentiated cells with Ezh2/CDK4/6 dual inhibitor compared to control across 3 MB cell lines without washout (n = 4 per cell line). P-values were calculated using a t-test. **D)** Number of differentiated cells following washout shows an increase in the total number of differentiated cells compared to cells that received only CDK4/6 inhibitor (n = 8). P-values calculated using ANOVA with post-hoc Tukey test.
